# Unprecedented Cytotoxicity of Transition Metal Metallacarboranes on Triple Negative Breast Cancer Cells and a Vertebrate Cancer Model

**DOI:** 10.1101/2025.01.29.635527

**Authors:** Neville Murphy, William J. Tipping, Yi Ding, Róisín M. Dwyer, Karen Faulds, Abhay Pandit, Duncan Graham, Herman P. Spaink, Pau Farràs

## Abstract

Metallacarboranes have long been the subject of attention in the context of medicinal chemistry because of their promising characteristics and unconventional interactions with biological entities. The metal centre has been shown to have a significant influence on the internalisation and cytotoxicity of compounds in human cell lines. Additionally, specific nanomolar concentrations of metallacarboranes have demonstrated toxic effects on MDA-MB-231 cells under in vitro and in vivo conditions.

**GRAPHICAL ABSTRACT:** 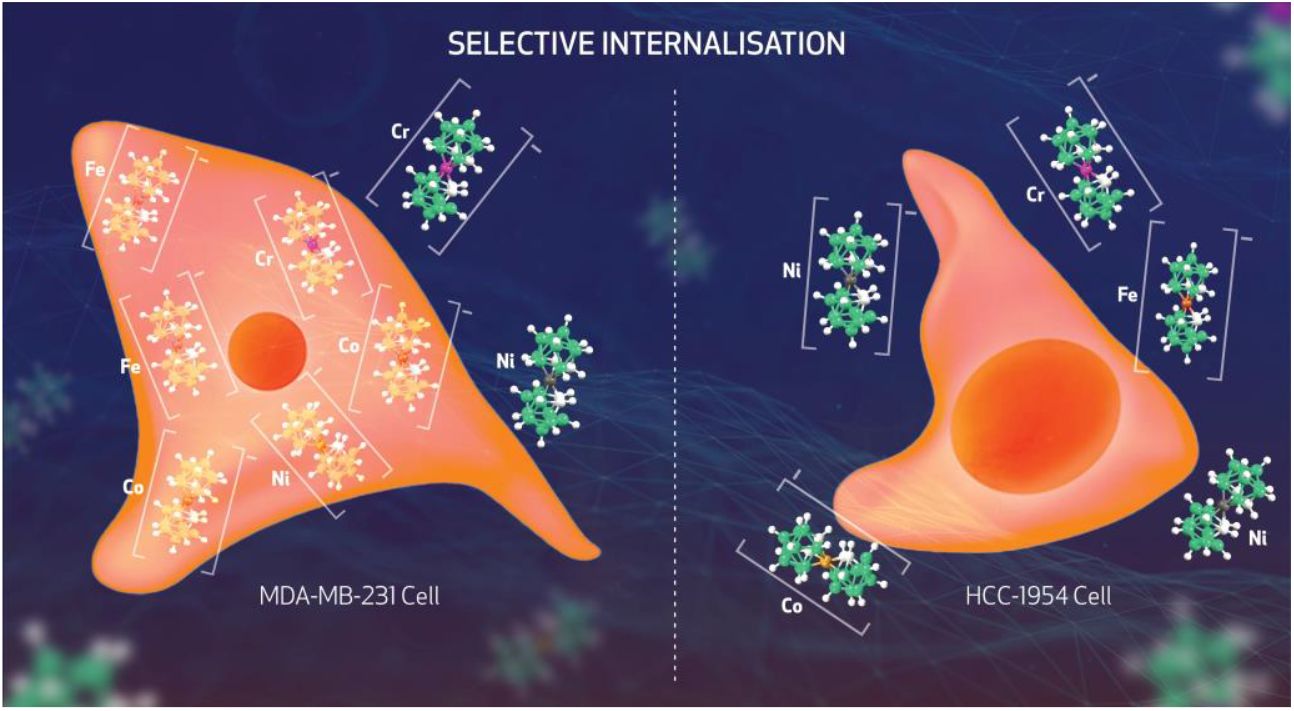

## INTRODUCTION

Triple-negative breast cancer (TNBC) is one of five subtypes of breast cancer. Occurring in 12-17% of female breast cancer patients, TNBC tumours lack expression of estrogen and progesterone receptors and lack amplification of the Her-2 protein. This lack of expression rules out a plethora of traditional therapeutics such as selective estrogen modulator receptors, and the monoclonal antibody trastuzumab as being effective treatments.^[1]^ The anthracyclines and taxanes are initially effective but are plagued with issues of resistance development, and return of the cancer post treatment.^[2,3]^ Application of a new class of therapeutics, with unique mechanisms of action could be hugely beneficial in this field, which has led many researchers to turn to boron clusters as a fresh approach to drug design.^[4–6]^ Boron itself has been successfully used in drug design with bortezumab, ixazomib (multiple myeloma) and tavaborole (onchocomytosis) being prominent examples.^[7]^ However, boron in cluster form has rarely been incorporated into commercial/natural product therapeutics with the exception of boron neutron capture therapy compounds such as sodium mercapto-undecahydro-closo-dodecaborate (BSH). This, coupled with the absence of boron clusters in nature may make these compounds less susceptible to mechanisms of resistance, offering a unique opportunity in this regard.^[8]^ Factors such as the carborane cage structure not having been metabolized before by biological systems and the unique hydrogen and dihydrogen interactions of the boron cluster cage hydrogens, are key to this theory.^[8]^

Metallacarboranes are a family of boron clusters that comprise of boron, hydrogen, carbon and a metal centre.^[9]^ Properties such as unconventional biological interactions, high stability and relatively low cytotoxicity typically associated with boron clusters has made them primary candidates for biomedical applications in recent times.^[4–6,10–12]^ The focus of this work is primarily the properties of the 3,3′-metalla-*commo*-bis(1,2-dicarba-*closo*-dodecaborane) class of metallacarboranes; a metal centre with two [C_2_B_9_H_11_]^2−^ ‘dicarbollide’ ligands either side. These compounds are often referred to with the elemental symbol followed by ‘SAN’, referring to the sandwich complex nature of the compounds. The cobalt bis-dicarbollide (CoSAN) has had the most attention by far, with extensive studies in the literature of its properties and potential applications.^[4,11,13–16]^ While not as much detail is available for other transition metal derivatives, there is enough data to show that there are significant differences in the properties of these compounds due to the metal centre. One such example is their redox potentials (**Table 1**).^[17]^ Involvement of transition metal-containing therapeutics in cellular processes has been reported previously as a valuable target in drug development.^[18]^, for example in the case of the iron bis-dicarbollide (FeSAN) which has promoted generation of reactive oxygen species (ROS) at low concentrations.^[19]^

**Table 1:** Redox potentials for Co, Fe and Ni metallacarboranes (in volts vs Fc^+^ /Fc). * Denotes a chemically irreversible redox pair.^[20]^

| M | M<sup>IV/III</sup> | M<sup>III/II</sup> | M<sup>II/I</sup> |
| --- | --- | --- | --- |
| Co | +1.22 | -1.75 | -2.64 |
| Fe | +0.76* | -0.78 |  |
| Ni | -0.17 | -1.01 | -2.52 |

Metallacarboranes however, have other potential avenues of cytotoxicity towards biological targets; the inhibition of HIV protease and carbonic anhydrase IX by derivatives of CoSAN serve as prominent examples of this.^[21,22]^ More generally, their ability to self-aggregate as well as having strong affinity for bovine serum albumin proteins demonstrate their potential as highly bioavailable entities in vivo, which have previously been shown to readily cross membranes.^[13,14,22–25]^ A key component of CoSAN’s success in this regard is thought to be the capability of the cage hydrogens to interact with proteins.^[26– 28]^ These can be quite diverse; hydrophobic, hydrophilic, and dihydrogen type-interactions have all been reported in protein-boron cluster pairs.^[28,29]^ These interactions have been linked to the partial atomic charges of the cage, which have been shown to be affected by the metal centre (**Table 2**). The more negative the partial atomic charge, the stronger the expected interactions with proton donor hydrogens in protein targets.^[30]^ The effect of these parameters on the internalisation and cytotoxicity of the metallacarboranes in the context of breast cancer remains relatively unknown. A study comparing cytotoxicity and internalisation of the cobalt and iron metallacarboranes in *C. elegans* was carried out recently with the use of EDX, XPS and NMR.^[31]^ It was found in this study that the iron was retained much better within the embryos than the cobalt, already highlighting the differences in behaviour between metal centres. However, more metal centres would need to be tested to establish trends, and monitoring of uptake in live subjects is preferable in drug localisation studies.

**Table 2:** Partial atomic charges of metallacarboranes obtained using the RESP methodology as seen in reference 32, which helps to describe the differing character of each cage hydrogen. Numbering (right) is repeated on the second cage in parentheses.

| Atom | Fe<sup>3+</sup> | Co<sup>3+</sup> | Ni<sup>3+</sup> |
| --- | --- | --- | --- |
| **M (3)** | 0.03 | 0.06 | 0.07 |
| **C (1,2)** | 0.02 | 0.02 | 0.01 |
| **B (4,7)** | 0.01 | 0.02 | 0.02 |
| **B (8)** | 0.01 | 0.01 | 0.0 |
| **B (5,11)** | 0.04 | 0.04 | 0.05 |
| **B (6)** | 0.01 | 0.0 | 0.0 |
| **B (9,12)** | 0.03 | 0.03 | 0.0 |
| **B (10)** | 0.06 | 0.07 | 0.07 |
| **H (1,2)** | 0.1 | 0.1 | 0.1 |
| **H (4,7)** | -0.1 | -0.1 | -0.1 |
| **H (8)** | -0.12 | -0.147 | -0.13 |
| **H (5,11)** | -0.1 | -0.11 | -0.1 |
| **H (6)** | -0.09 | -0.09 | -0.09 |
| **H (9,12)** | -0.13 | -0.13 | -0.12 |
| **H (10)** | -0.12 | -0.12 | -0.12 |

To this end, fluorescence has long been the primary strategy for cellular imaging, enabling the mapping of cellular structures and tracking of tagged compounds within them.^[32]^ Metallacarboranes themselves have been successfully tracked within HEK293 cells using a fluorescent tag.^[33]^ For the purposes of this study however, the addition of bulky fluorescent tags would interfere with the desired analysis of the interactions of the cage of these compounds with the cellular environment. Stimulated Raman scattering microscopy (SRS) has provided label-free bioorthogonal imaging of cells, while also tracking compounds containing Raman-active vibrational bands within the cell-silent window.^[32,34]^ While spontaneous Raman has been previously applied to image metallacarboranes within cells, the relatively low sensitivity of the technique requires concentrations within the mM range.^[35]^ SRS imaging enables higher sensitivity detection compared to spontaneous Raman imaging. Therefore, the detection of drugs and small molecules using concentrations closer to the IC_50_ can be achieved to provide a clearer understanding of cellular localisation and concentration.

This work reports the changes in metallacarborane behaviour brought about by various metal centres in the context of breast cancer, namely Co, Fe, Cr and Ni, which provide a palette of d electron counts and varying electronegativities. While there has been significant interest in the application of boron clusters and metallacarboranes in the field of cancer, few have addressed breast cancer specifically, with even fewer examples for the full sandwich metallacarboranes.^[4,6]^ One recent example of such a study is presented in the work of Bednarska-Szczepaniak *et al*., testing Co, Fe and Cr metallacarboranes on ovarian cancer cell lines.^[36]^ It was found that metallacarborane internalisation could be linked to a marked increase in mitochondrial activity, ‘exhausting’ the cell which lead to cell death. Cell death due to exhaustion was especially prevalent in the iron metallacarborane. This work evaluated the cytotoxicity of the selected compounds using colorimetric cell viability assays in HER2 amplified HCC-1954 and the triple-negative MDA-MB-231 breast cancer cell lines. This enabled a direct comparison between breast cancer types and normal human dermal fibroblast (HDF) cell lines to provide cytotoxicity data on a non-tumourigenic cell line.

Some metallacarboranes exhibited high potency against triple-negative MDA-MB-231 cells with negligible cytotoxic effects on other cell lines. Cellular uptake was monitored using stimulated Raman scattering microscopy in MDA-MB-231 and HCC-1954 cell lines, which matched the trends observed in cytotoxicity. Metallacarboranes were also found to have significant cytotoxic effects at nanomolar concentrations compared to MDA-MB-231 cells in vitro and in vivo. Herein, we present the first comprehensive set of data on various metallacarboranes exhibiting selective cytotoxicity against human cancer cell lines, with exciting new insights into their potential as therapeutic agents.

## RESULTS

### Metallacarboranes exhibit selective cytoxicity vs TNBC cell lines

MDA-MB-231 is a TNBC cell line isolated from the pleural effusion of a patient with invasive ductal carcinoma which has been classified as basal, as it lacks the expression of estrogen receptor (ER), progesterone receptor, and human epidermal growth factor receptor 2 (HER2). Suppressed expression of claudin-3 and claudin-4 has been classified as the claudin-low molecular subtype. Its aggressive and invasive nature has made it a useful model of late stage breast cancer.^[37]^ Therefore, cell viability tests following treatment with various dicarbollides would provide information on the suitability of each metal centre for TNBC therapeutics. Metallacarboranes and paclitaxel (an established therapeutic in breast cancer treatment) were tested at concentrations ranging from 100 down to 6.25 µM for 72 h (**Figure 1A**). FeSAN showed the most promise with mean cell viability remaining below 50% up until 12.5 µM, and significant reduction compared to control at 6.25 µM. CoSAN was the next most effective, with mean cell viability rising above 50% at 25 µM, and a significant reduction compared to the control up until 12.5 µM. Interestingly, treatment with CrSAN and NiSAN did not significantly affect cell viability below 50 µM concentrations, demonstrating selectivity between compounds on the MDA-MB-231 cell line. This may arise from the differing properties of the cage hydrogens and their interactions with cellular targets, as well as redox properties brought about by the metal centre within the compound. Paclitaxel had a significant effect on cell viability until 25 µM but was outperformed by FeSAN in terms of reducing cell viability following 72 h treatment.

**Figure 1:**
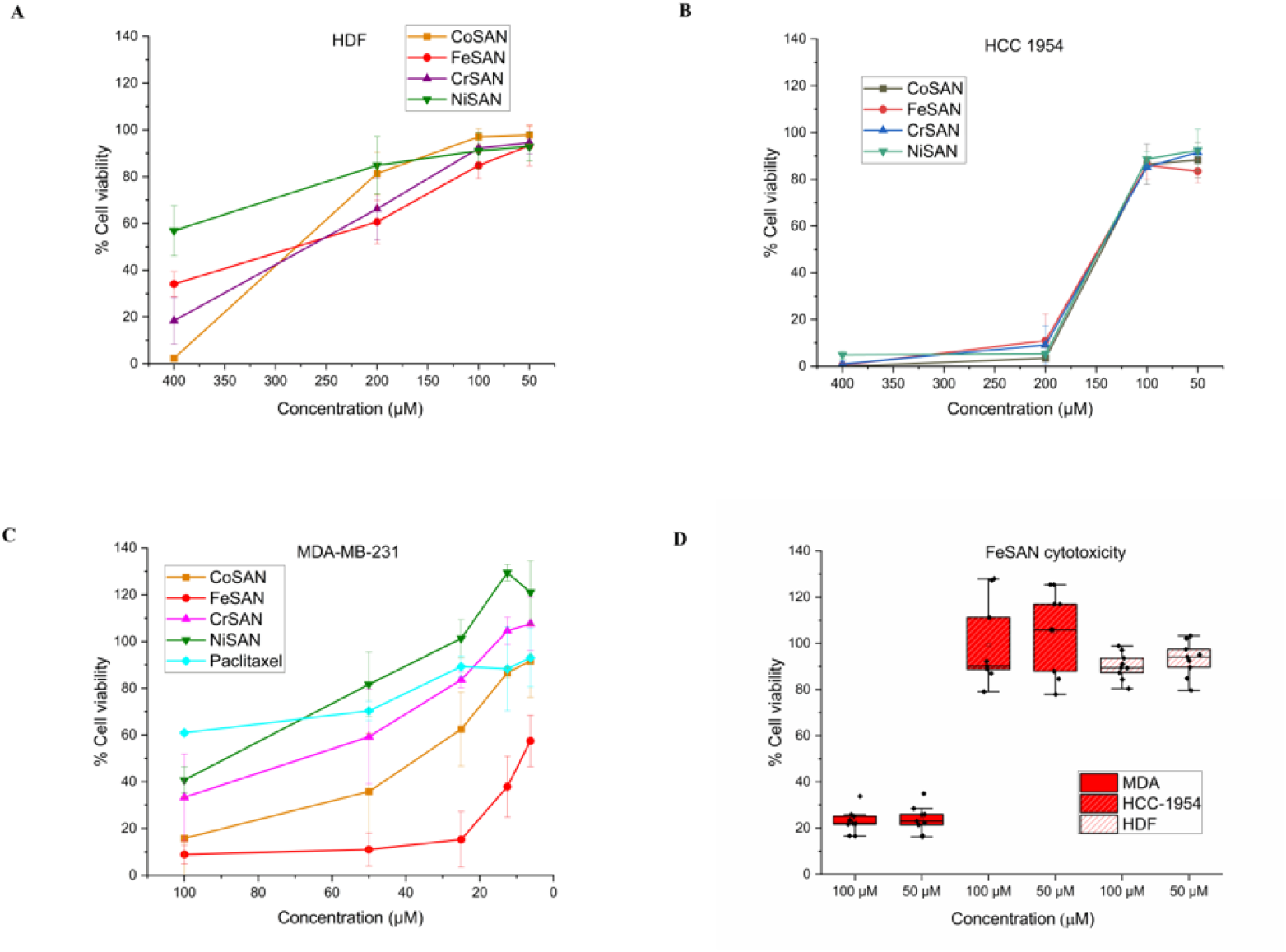
Cell viability of **A**: HDF, **B**: HCC-1954 and **C**: MDA-MB-231 cell lines following 72 h incubation with metallacarboranes at stated concentrations. **D**: Selectivity between cell lines in FeSAN shown with box plot. This data highlights the selectivity of the metallacarboranes cytotoxicity towards the TNBC MDA-MB-231 cell line, as well as the higher potency of FeSAN.

The effects of Co, Fe, Cr, and Ni metallacarboranes on cell viability were carried out from 400 to 50 µM at 24 h (**Figure S1**). It was found that the cytotoxicity of the compounds was generally too low to be seriously considered potent, with cell viability well above 50% at 100 µM. The significant increase in cytotoxicity observed across all compounds with a treatment time of 72 h indicates that toxicity is dependent on exposure time and concentration.

To determine selectivity between breast cancer cell lines, HER2 positive HCC-1954 cells were treated with metallacarboranes at concentrations 400 µM to 50 µM (**Figure 1B**). This cell line was far more resilient to the cytotoxic effects of the metallacarboranes; at 100 µM across all metal centres the cell viability is much higher, approaching 100% and displaying no significant change in effect compared to control. Combined, these results indicated that the cytotoxicity observed on the MDA-MB-231 cell line is selective.

Human dermal fibroblasts (HDF) are a normal human cell line that originate from mesenchymal cells. They have previously featured in cytotoxicity studies with MDA-MB-231 cell lines as a comparative measurement.^[38,39]^ Their role in this study was to determine if there was selectivity in the effect on cell viability between a triple negative breast cancer cell line (MDA-MB-231) and a non-invasive, slower metabolising normal human cell line. The results for 72 h incubation at 400 µM to 50 µM concentrations are presented in **Figure 1C**. Metallacarboranes were shown to only have a significant effect on cytotoxicity at 400 µM, with partial inhibition of cell viability visible at 200 µM, although the effect was not significant. Compared to the relatively high cytotoxicity observed at 100 µM in the MDA-MB-231 cell line, this data represents another example of selectivity between cell lines in the metallacarboranes. The significant differences in cytotoxic effects between cell lines as well as metal centres may be ascribed to factors such as varying levels of cellular internalization and affinity for cellular protein targets, which will require further studies.

### Metallacarborane cellular uptake is linked to cytotoxicity

Before discussion of specific cellular targets, one potential explanation for the selectivity observed with cytotoxicity between the metal centres and cell lines is the uptake. The B–H stretching vibrations from boron clusters are represented by a broad peak at 2480-2680 cm^-1^ in the Raman spectrum. Metallacarboranes have been previously detected within human HEK293 cells using spontaneous Raman spectroscopy.^37,42^ As shown in our previous work, metallacarboranes may be effectively used in multiplex detection using SRS at much lower concentrations, while providing higher spatial resolution.^43^ Following single acquisitions of the Raman spectra of the metallacarboranes as solids (**Figure S2**), spontaneous Raman scattering mapping of each metallacarborane species was then carried out on MDA-MB-231 and HCC-1954 cell lines. Each cell line was treated with metallacarborane (500 µM, 30 min) representing a shorter incubation time and lower concentration compared to previous Raman-based studies, owing to the observed cytotoxic effects with longer exposure times.

While the compounds were detected, the signal was too weak to make conclusive comparisons of cellular B–H Raman intensity between compounds (**Figure S3, S4)**. To effectively compare metallacarborane species uptake, HCC-1954 and MDA-MB-231 cell lines were seeded onto glass slides and subjected to 500 µM of the metallacarborane in question for 15 min, then SRS intensity was acquired at 2930 cm^-1^ (CH_3_, proteins, symmetric stretch) and 2851 cm^-1^ (CH_2_, lipids, symmetric stretch) to determine the cells as regions of interest. SRS intensity was then measured at ∼2570 cm^-1^ to determine the relative uptake of the compounds, and finally at 2400 cm^-1^ to measure the background off-resonance signal. We quantified the B–H signal in individual cells to enable a comparison of metallacarborane uptake as a function of metal centre. The degree of metallacarborane uptake appeared to be governed by the metal centre and the cell line in question (**Figure 2A**). In MDA-MB-231 cells, FeSAN showed the most effective uptake, followed by NiSAN and CoSAN. CrSAN was the least effective, although it still demonstrated significantly more SRS signals than the control **(Figure 2B**). These data demonstrate the capability of all tested metal centres to internalise within MDA-MB-231 cells in a relatively short period. The HCC-1954 cell line showed a slight increase in SRS intensity with 500 µM metallacarborane treatment, although the difference was not statistically significant (**Figure 2B**). This aligns well with the cytotoxicity data, in that the lower internalization of the compounds in the HCC-1954 cell line coincides with the relatively low cytotoxicity of metallacarboranes.

**Figure 2:**
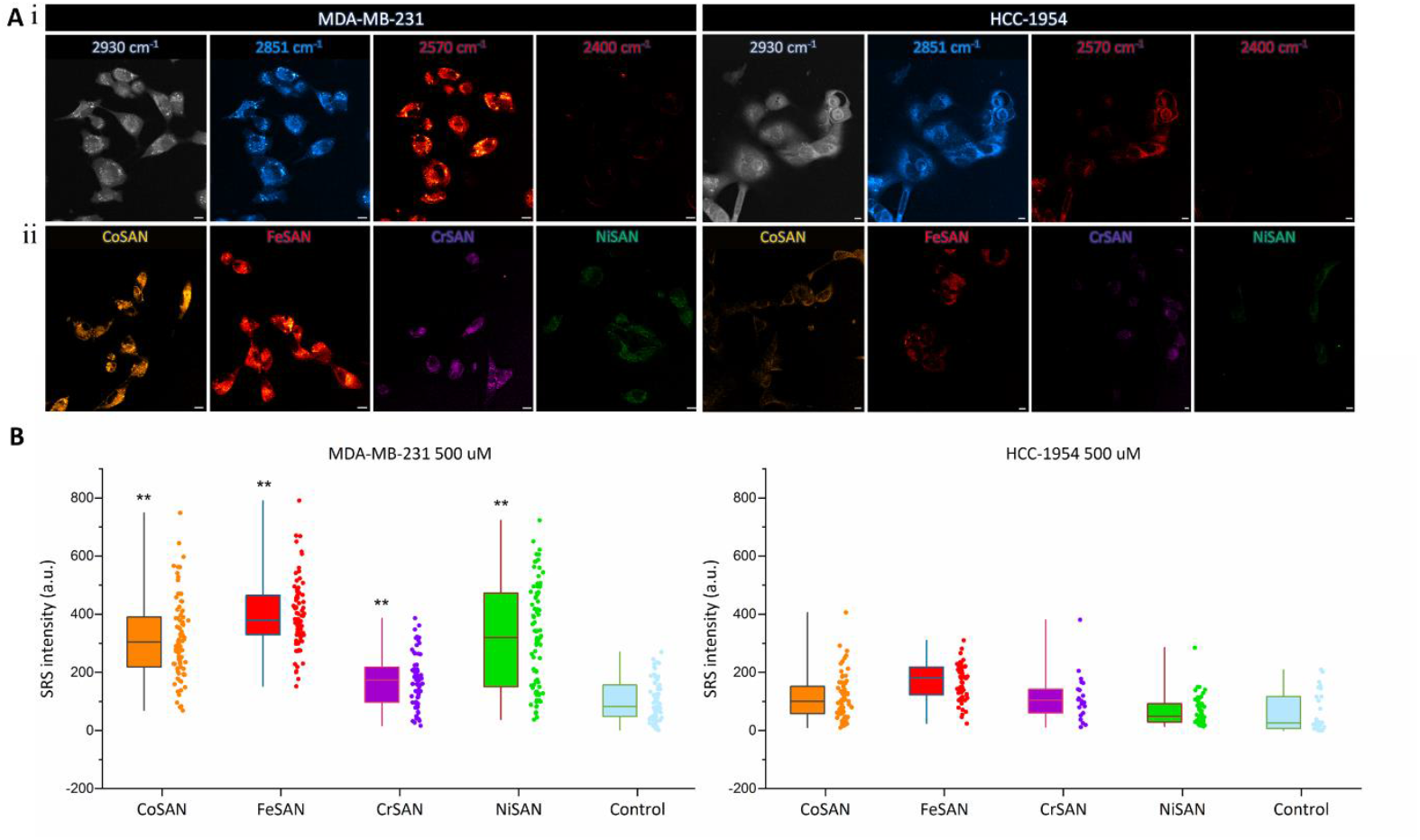
**A**: (i) SRS imaging of MDA-MB-231 (left) and HCC-1954 (right) cell lines at 2930 cm^-1^ (CH_3_, proteins), 2851 cm^-1^ (CH_2_, lipids), 2570 cm^-1^ (B–H, FeSAN) and 2400 cm^-1^ (off-resonance). Look-up tables: 2930 cm^-1^ (greyscale, 0–3000 a.u.), 2851 cm^-1^ (cyan hot, 0–3000 a.u.) 2570 cm^-1^ (red hot, 0– 2500 a.u.), 2400 cm^-1^ (red hot, 0–2500 a.u.). Scale bars: 10 μm. (ii) SRS images at 2570 cm^-1^ which have been background subtracted using the images acquired at 2400 cm^-1^. Look-up tables: CoSAN (orange hot, 0–2500 a.u.), FeSAN (red hot, 0–2500 a.u.) CrSAN (magenta hot, 0–2500 a.u.), NiSAN (green, 0–2500 a.u.). Scale bars: 10 μm. **B**: SRS intensity values from within MDA MDA-MB-231 (left) and HCC-1954 (right) cell lines following incubation with metallacarboranes, detailing much higher uptake in the mDA-MB-231 cell lines compared to the HCC-1954. Data extracted through ImageJ software, using 2930 cm^-1^ image as a mask, then setting measurements to the images at 2570 cm^-1^ which had been background subtracted using the images acquired at 2400 cm^-1^. *P ≤ 0.05, **P ≤ 0.01, ***P ≤ 0.001 (student’s t-test).

Cellular localisation of the metallacarboranes has been previously shown to be primarily in the cytoplasm in a selection of human cell lines, though none on human breast cancer cell lines.^[35,40,41]^ Comparing the SRS intensity of the B–H stretch at 2570 cm^-1^ within the nucleus and the cytoplasm of the MDA-MB-231 cells following treatment with the metallacarboranes presented another example of this phenomenon. Interestingly, the trends observed regarding SRS intensity increases between metal centres are similar between the nucleus and cytoplasm, with nuclear SRS intensity being significantly higher than the control in all metallacarborane species (**Figure 3A**). This implies a high degree of nuclear penetration in the MDA-MB-231 cells. Further studies using perfusion chambers to achieve imaging in situ upon treatment of cells with metallacarboranes showed detectable internalisation of FeSAN after 2 minutes (**Figure S5**) using infusions of 500 and 250 µM concentrations (**Figure 3B**). Interestingly, there were no significant differences in SRS intensity between 500 and 250 µM (**Figure 3B**). Interestingly, there were no significant differences in the SRS intensity between 500 and 250 µM after two minutes (**Figure S6**), indicating that 250 µM may be the saturation point for the uptake phenomenon. CrSAN, which had the lowest mean SRS intensity following treatment, was also tested for comparison and showed no discernible change in intensity after 2 minutes incubation. The weaker signal may be related to the slower internalization of CrSAN compared to FeSAN (**Figure S7**). While FeSAN and NiSAN were internalized most effectively, showing the highest mean SRS intensities, FeSAN was far more toxic to cells. Our results indicated rapid cytoplasmic internalisation of all tested metallacarboranes (with the exception of CrSAN) in MDA-MB-231 cells, and we are further investigating the source of the selectivity and mechanism of internalisation.

**Figure 3:**
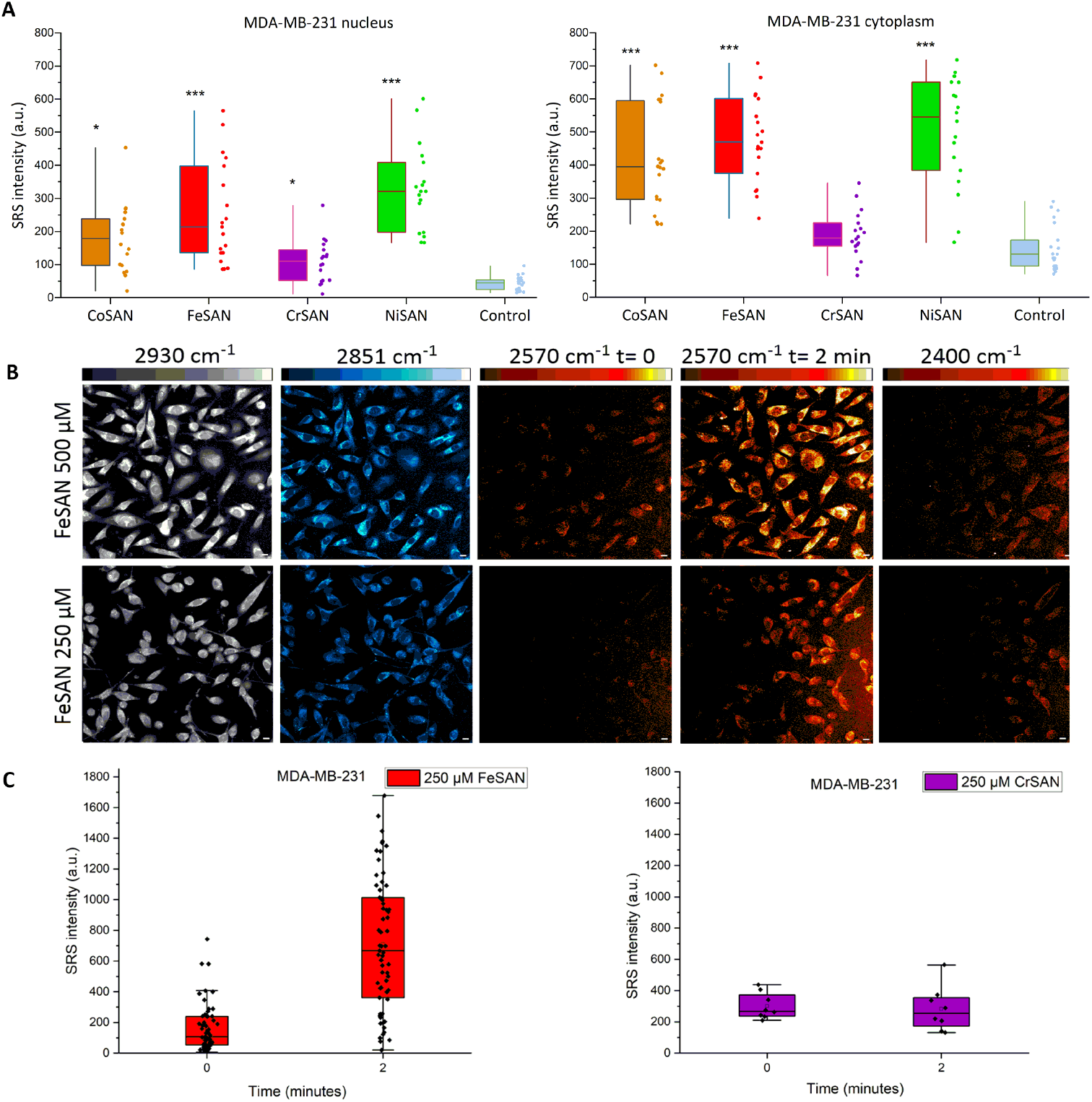
**A**: SRS intensity values from within MDA MDA-MB-231 cells following incubation with metallacarboranes in the nucleus (left) and cytoplasm (right), showing significant SRS intensity increases in both, but to a lesser extent in the nucleus. Data extracted through ImageJ software, using 2930 cm^-1^ image as a mask, then setting measurements to the images at 2570 cm^-1^ which had been background subtracted using the images acquired at 2400 cm^-1^. The area of interest was then established manually using imageJ drawing tool. **B**: SRS imaging of MDA-MB-231 cells at 2930 cm^-1^ (CH_3_, proteins), 2851 cm^-1^ (CH_2_, lipids), 2570 cm^-1^ (B–H) at t= 0 min (pre-treatment), t= 2 min and 2400 cm^-1^ (off-resonance) visually demonstrating speed of uptake of the compounds. Look-up tables: 2930 cm^-1^ (greyscale, 0–3000 a.u.), 2851 cm^-1^ (cyan hot, 0–3000 a.u.) 2570 cm^-1^ (red hot, 0–2500 a.u.), 2400 cm^-1^ (red hot, 0–2500 a.u.). Scale bars: 10 μm. **C**: SRS intensity at t= 0 min (pre-treatment), t= 2 min post treatment for FeSAN (left) and CrSAN) right, highlighting selectivity n uptake between FeSAN and CrSAN in MDA-MB-231 cells. Data extracted through ImageJ software, using 2930 cm^-1^ image as a mask, then setting measurements to the images at 2570 cm^-1^ which had been background subtracted using the images acquired at 2400 cm^-1^. *P ≤ 0.05, **P ≤ 0.01, ***P ≤ 0.001 (student’s t-test).

Therefore, the lower observed cytotoxicity of CrSAN may be attributed to the poorer uptake of the compound due to the character of the cage hydrogens, which have been shown to play an important role in membrane crossing.^[42]^ Interestingly, NiSAN demonstrated a high degree of internalisation, comparable to that of FeSAN, despite having minimal effect on the cell viability in comparison to the other metal centres. This may be attributed to the fact that NiSAN begins to switch to the insoluble Ni(IV) salt over time in PBS solution, (**Figure S8**), which may reduce its potency in a 72 h treatment period, and that changes in oxidation state have a direct effect on the electron density in the metallacarborane cage, affecting its capacity to interact with neighbouring species.^[10]^ NiSAN which had been incubated at 37 C° in cell culture media for 48 h, thereby resulting in a reduction in the available Ni(iii) in the culture media. This was tested for internalisation in MDA-MB-231 cells using SRS spectroscopy. No significant increase in the SRS intensity was detected, confirming our hypothesis regarding the lack of cytotoxicity (**Figure. S9**). It can also be hypothesised that the effects of the metal centre on the cage hydrogens govern the ability of the compounds to internalise within MDA-MB-231, thus reflecting upon the overall cytotoxicity observed. However, other effects, such as binding to potential targets within the cell, could be factors governing the cellular uptake of metallacarboranes.

### Metallacarboranes exhibit cytotoxicity vs TNBC cell lines at specific nanomolar concentrations

In the literature, there is precedent for changes in the behaviour of metallacarboranes at nanomolar concentrations. Kaniowski *et al*. reported an increase in reactive oxygen species with both FeSAN and anti-sense nucleotide derivatives of FeSAN in HeLa cells.^[19]^ An increase in metabolic activity was observed in HeLa cells at 25 nM concentration, which was attributed to the redox activity of the iron metal centre generating ROS and thus activating the defence mechanisms of the cells. To determine the effect of different metal centres and therefore a library of redox activities, the four transition metal metallacarboranes (**Figure 3A**) were tested against MDA-MB-231 cells in a similar range (20-2.5 nM) for effects on cell viability. Interestingly, a significant change in cell viability compared to the control was observed at 5 nM CoSAN, FeSAN, and CrSAN, which tapered off with increasing and decreasing concentrations. The effect was especially significant in FeSAN, and less so for CoSAN, CrSAN mirroring behaviour was observed at micromolar concentrations. NiSAN induced a significant increase in cell viability at 20 and 1.25 nM, despite the decrease in cell viability observed at 5 nM. This represents a very different response of the cells to the HeLa study in the literature, where viability did not drop below control in the case of FeSAN.^[19]^ To investigate if this phenomenon may be explained by the effects seen in the literature with the increase of ROS generation in HeLa cells following treatment with FeSAN, a 2’,7’-dichlorofluorescein diacetate/2’,7’-dichlorodihydrofluorescein diacetate DCFDA/H_2_DCFDA assay was carried out which detects ROS, with *tert*-butyl hydroperoxide (TBHP), a known ROS generator as positive control which was then used to normalise other results and cells without any treatment as negative control (**Figure 4A**). No significant ROS generation was observed, which revealed no significant production of ROS compared to control, though an increase was observed in the case of NiSAN, which had increased cell viability. Slight increases in ROS have previously been shown to increase the metabolic activity of cells.^[19]^ This indicates a different mechanism of action of the compounds at this concentration in the MDA-MB-231 cell line compared to the activity reported for the HeLa cell line in the literature.

**Figure 4:**
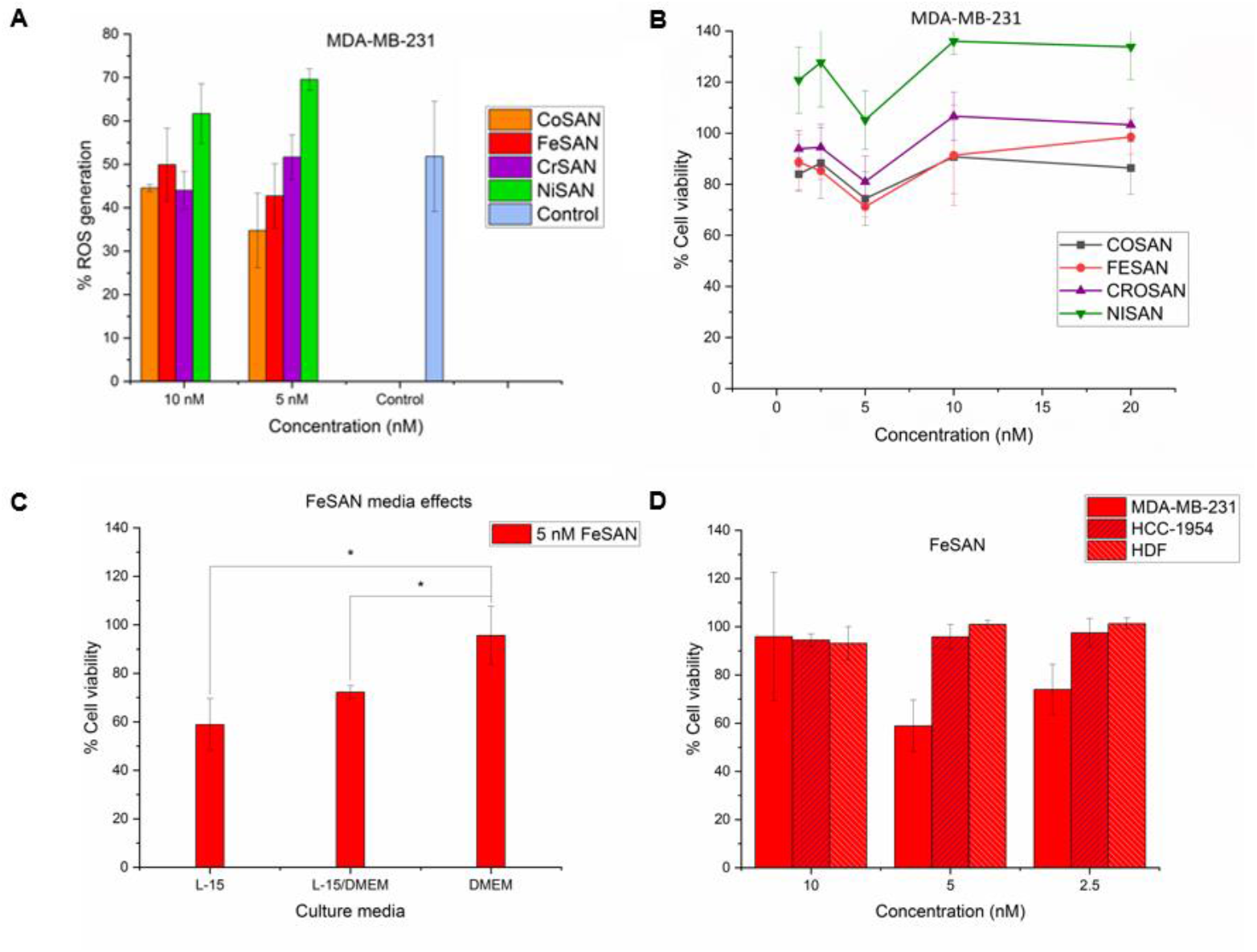
Cell viability and ROS production of metallacarboranes at nanomolar concentrations. **A**: Results of DCFDA assay for detection of ROS in MDA-MB-231 cells showing increase in ROS in presence of NiSAN. Data collected following metallacarborane treatment normalised against cells treated with TBHP, a known ROS producer. **B:** Cell viability normalised vs. control following treatment of MDA-MB-231 cells cultured in L-15 with metallacarboranes, showing the drop in cell viability at 5 nM. **C**: Viability of MDA-MB-231 cells following treatment with FeSAN in culture and assayed in L-15, cultured in DMEM and assayed in L-15, and finally cultured and assayed in DMEM, displaying selectivity of the effect with regard t cell culture conditions. **D:** Cell Viability data of MDA-MB-231, HCC-1954, and HDF cell lines following treatment with nM concentrations of FeSAN for 72 h. *P ≤ 0.05 **P ≤ 0.01 ***P ≤ 0.001 (student’s t-test). These results once again hihghlighting selectivity, this time between cell line types.

The media in cell culture has been previously shown to have significant effects on the genome of cells, with one case study on MDA-MB-231 cells detecting over 9,000 changes in the genome following use of different media.^[43]^. DMEM (glucose) and L-15 (galactose) media contained differing amounts of sugars and amino acid supplements. Changes in these parameters have been shown to trigger a range of genetic and metabolic changes to the cells, which would provide some indication as to whether the toxicity is general or more targeted in character.^[44]^ It was indeed found that the culture conditions of the cells have an influence over these cytotoxic effects. After culturing for two months in DMEM, the cells were tested in the same concentration range in the presence of FeSAN for 72 h, and the results are presented in **Figure 4B**. Cells cultured in DMEM were shown to be more resilient to nanomolar FeSAN exposure at nM concentrations, with no significant reduction in cell viability. To ascertain whether the origin was due to interactions of FeSAN with the media during the incubation period, the cells were cultured in L-15 media, and the assay was carried out in DMEM. The cell viability was still reduced by ∼30%, significant to control and DMEM cultured and assayed cells, indicating that the effects seen on L-15 cultured cells may be due to the expression of certain targets being affected by the supplements in the cell culture media. HCC-1954 and HDF cell lines were also subjected to 10 to 2.5 nM of FeSAN for 72 h to check for similar effects and monitor the selectivity of the phenomenon, with the HDF cell lines in DMEM and the HCC-1954 cell line in RPMI (**Figure 4D**). No significant cytotoxicity was observed in other cell lines, indicating selectivity for cytotoxicity. SRS intensities at 2570 cm^-1^ were measured within the cells after 72 h of incubation at 25, 10, and 5 nM. Although the average SRS intensity was higher for the 5 nM incubation, it was not statistically significant (**Figure S9**). It is possible that the mean increase in intensity could be due to the aggregation of the metallacarboranes at 5 nM concentration being favourable for accumulation within the cells, which may also explain the observed cytotoxicity. However, this topic requires further analysis before more concrete conclusions can be drawn regarding the source of cytotoxicity.

### In vivo cytotoxicity observed within zebrafish MDA-MB-231 model

Having determined cytotoxicity at low micromolar values, as well as nanomolar concentrations, an alternative, physiologically relevant model was explored to test the capabilities of the compounds. Zebrafish are an established model for drug discovery in cancer, due to transparency which enables relatively straightforward imaging, and a similar genetic make up to humans which allows study of metastasis and cancer development in a relevant environment.^[45,46]^ First, the maximum tolerable concentrations were determined using GFP-labelled vasculature.

Fli:GFP casper zebrafish larvae (72 h.p.f.). Compounds at different concentrations were introduced into the egg water of 2 day old, hatched zebrafish larvae. The survival rate was then assessed five days post fertilization (d.p.f.). The results showed that only nanomolar concentrations were within the therapeutic window, with the exception at 5 nM in the case of CoSAN (**Table 3**).

**Table 3:** Survival rates of 5 d.p.f. zebrafish larvae found to be unaffected at 1 µM and lower with Co, Cr and Ni metallacarboranes; with the exception of toxic effects at 5 nM in the case of CoSAN. Fe metallacarborane was found to be potentially therapeutic to the zebrafish larvae only at 5 nM concentration.

| Concentration<br>(μM) | CoSAN | FeSAN | CrSAN | NiSAN |
| --- | --- | --- | --- | --- |
|  | % Live fish |  |  |  |
| 10 | 0 | 0 | 0 | 0 |
| 1 | 100 | 0 | 100 | 100 |
| 0.1 | 100 | 83 | 100 | 100 |
| 0.005 | 83 | 100 | 100 | 100 |

This is another example of abnormal changes in the biological activity of the compounds at this concentration, Zebrafish were completely intolerant to the compounds at micromolar concentrations, even at higher nanomolar concentrations, in the case of FeSAN (**Table 3**). However, the compounds showed more promise at 5 nM, with CoSAN being the exception; among the 83% surviving fish, some had physical defects. No cytotoxicity was observed under other conditions in the 18 larvae.

To monitor the effects of metallacarboranes on cells in vivo, CM-Dil stained MDA-MD-231 cells were prepared and injected into the avascular structure perivitelline space of 48 h.p.f. zebrafish larvae using an automated injection robot (Life Science Methods. B.V.).^[47]^ Cm-Dil staining has been successfully used previously to track cancer cell migration and proliferation, as the fluorescent dye intercalates with the cell membrane facilitating tracking.^[48]^ The larvae were then imaged using fluorescence microscopy on the whole fish at 1.5x and the injection site at 4x magnification, then treated with stated conditions for 72 hours. They were then imaged again, and the fluorescence intensity was compared to determine the cytotoxic effects of the treatment on the cancer cells **(Figure 5A)**.

**Figure 5:**
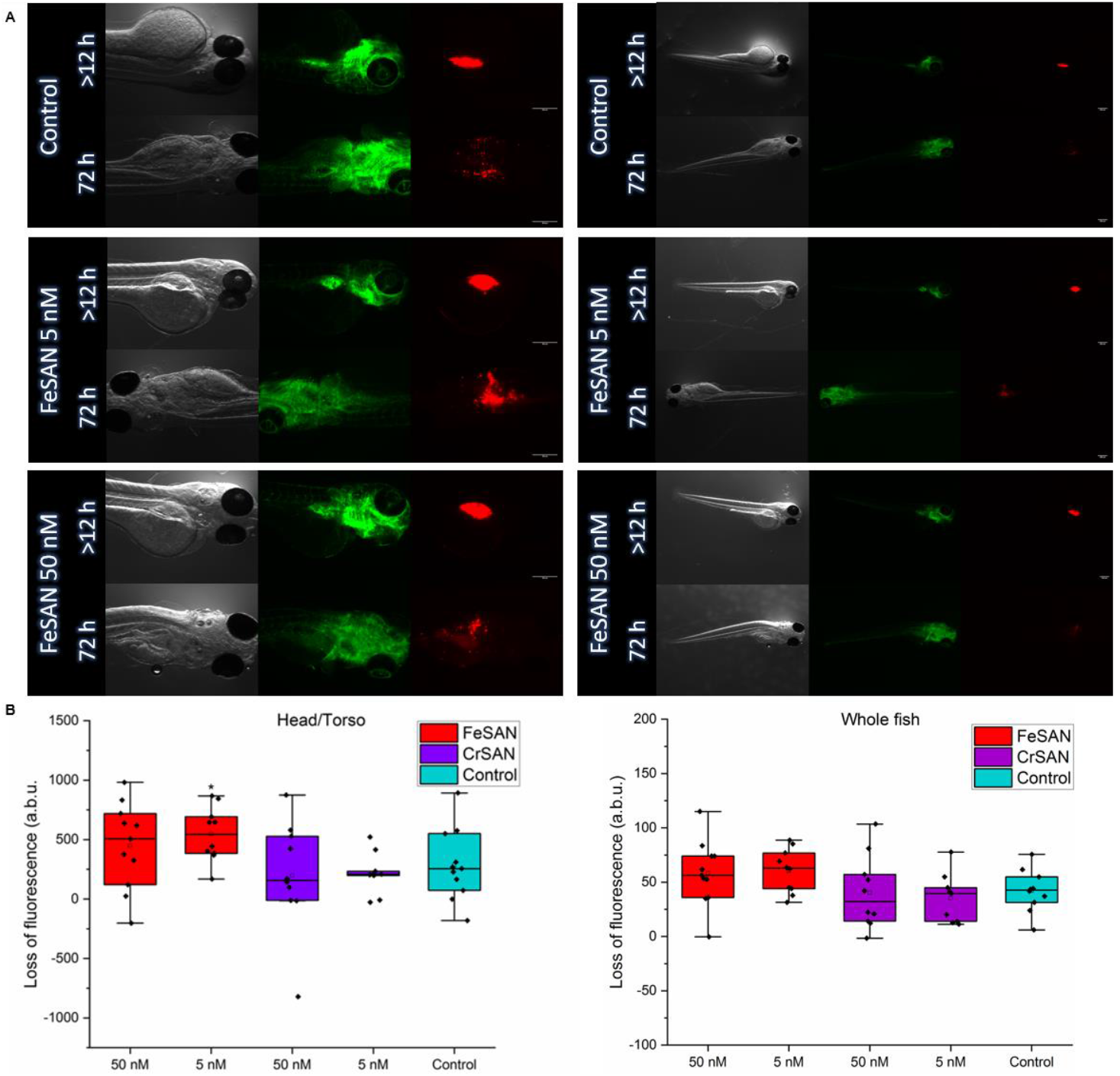
Details of the effect of metallacarboranes on CM-Dil-stained MDA-MB-231 cells injected into FLI:GFP Casper zebrafish. **A**: Images of fluorescence detected in FLI:GFP Casper zebrafish larvae post-injection at 48 h.p.f. and post-treatment with metallacarboranes vs. controls (120 h.p.i.). **B:** Calculated loss of fluorescence between injection at 48 h.p.i. and post-treatment with metallacarboranes versus controls (120 h.p.i.). Data are means of values for at least nine fish under each condition. Left: fluorescence measured at x4 magnification at the injection site in the perivitilline space, right: x1.5 magnification of the entire fish larvae. *P ≤ 0.05 (Student’s t-test).

FeSAN and CrSAN exhibited significant cytotoxicity to the MDA-MB-231 cell line in vivo at 5 nM, but did not have any effect on the larvae. Given the toxic effects on the larvae exhibited by CoSAN, and a lack of effect on the MDA-MB-231 cells by NiSAN, CrSAN, and FeSAN, they were chosen for the following in vivo studies. Survival rates were shown to be dependent on the toleration of the injections rather than metallacarborane treatment; uninjected fish in all controls survived in the study.

Images at x1.5 magnification were analysed for significant fluorescence reduction compared with the control. While no significant differences were observed, a trend was observed in that the 5 nM treatment led to a higher mean change in fluorescence than the control fish (**Figure 5B**). Therefore, images at higher magnification (x4) centred on the injection site and perivitelline space were analysed (**Figure 5A**). Interestingly, differences in fluorescence indicated a statistically significant change in the case of the 5 nM treatment with FeSAN over 72 h compared to the control, which aligns well with the data gathered in vitro **(Figure 5B)**. While the mean values of the FeSAN 50 nM treatment were similar to those at 5 nM, the wider range of fluorescence loss in the data set failed to make a significant change compared to the control, further highlighting the importance of the 5 nM concentration for cytotoxicity. CrSAN treatments, however, did not trigger any significant differences in fluorescence readings compared to the control, following the observed trend at micromolar concentrations compared to FeSAN. Beyond further demonstration of the cytotoxicity of FeSAN at 5 nM, these data represent the activity of FeSAN vs. a triple-negative breast cancer cell line in vivo, within a therapeutic dose for the animal model, shedding new light on the effects of these compounds at nanomolar concentrations. This compound could be applied in adjuvant therapy for patients with triple-negative breast cancer.

However, the source of cytotoxicity remains unclear and requires further extensive studies. Certainly, testing in more mature zebrafish models as well as other animals such as mice which may tolerate higher concentrations of the compounds would provide useful information on the activity at low micromolar concentrations, as well as analysis of more cancer subtypes to determine if any further trends can be observed. Regarding the uptake, knock out studies on membrane transporters, studying trends in uptake by different cell types and structure-uptake studies could shed light on the mechanism of uptake in this context. While there is evidence in the literature of CoSAN crossing membranes via concentration gradients and a ‘‘cooperative flip-flop’’ mechanism, this does not explain the lack of uptake in HCC-1954 cells nor the reduced capabilities of the anionic CrSAN to internalise in both cell types.^[49]^

## DISCUSSION

Overall, this work demonstrates the diversity of biological properties within species of metallacarboranes in a range of cell lines, through their cytotoxicity, uptake, and selectivity. By changing the transition metal centre, the cytotoxicity, uptake, and selectivity have been shown to vary between metallacarboranes and the two breast cancer cell lines. Iron metallacarborane has been shown to be the most promising candidate for TNBC therapy.

Additionally, a correlation has been found between cellular internalization and the cytotoxicity of metallacarboranes in MDA-MB-231 cells, which was observed within seconds of exposure at low micromolar concentrations. The source of cytotoxicity observed at micromolar concentrations is also unknown, as is the mechanism of transport across the cell membrane of MDA-MB-231 cells. Studies of metallacarborane uptake in a variety of cell lines to determine trends in cellular uptake, such as lipid content and membrane transporters, are key to explaining the phenomena observed in this study. In addition, cytotoxicity at low nanomolar concentrations, which were not toxic to other cell lines or Fli:GFP casper zebrafish larvae was observed. These effects are not immediately attributable to redox activity as previously reported with other human cell lines. FeSAN has shown promise as a future therapeutic, displaying cytotoxicity versus MDA-MB-231 cell line in vivo at therapeutic concentrations. Further studies with all cell lines cultured in L-15 media would shed more light on the effects of L-15 culture on the observed cytotoxicity seen in the MDA-MB-231 cell line. If no cytotoxicity is observed even under these conditions, it may indicate the targets of the metallacarboranes are exclusive to the MDA-MB-231 cell line. Cytotoxicity would implicate the L-15 media triggers expression of targets in multiple cell lines. Short tandem repeat {STR) profiling to determine the influence of the cell culture media on the genome, and which changes may be triggering the loss of cytotoxicity could help identify targets. Studies to shed light on these phenomena are currently underway.

Described herein is the first evidence that metallacarborane cytotoxicity in MDA-MB-231 cells may be linked to rate and overall levels of internalization. This data further highlights the importance of the metal centre in the biological behaviour of metallacarboranes. This is also the first report of the phenomenon of metallacarborane cytotoxicity exclusively at a specific nanomolar concentration, which was demonstrated in vivo and in vitro. However, the zebrafish vertebrate model used in the study may be improved upon in terms of physiological relevance to the cancer microenvironment in humans. Use of older zebrafish or 9proceeding to larger animals such as mice would provide this. Additionally, more detailed studies on the mechanisms of metallacarborane cytotoxicity are required, with PCR profiling of cells following exposure and screening for cellular targets being examples of such studies. This work does provide a suitable steppingstone for determining future applications as a combinational therapy with established anti-cancer drugs for the metallacarboranes.

## RESOURCE AVAILABILITY

### Lead contact

Further information and requests for resources and reagents used in this study should be directed to the lead contact, Pau Farràs.

### Materials availability

Experimental materials generated in this study are available from the lead contact with a completed Materials Transfer Agreement.

### Data and code availability

The raw data files generated during this study are available from the corresponding author (PF) upon request.

## ACKNOWLEDGEMENTS

This publication has emanated from research supported in part by a research grant from Research Ireland and is co-funded under the European Regional Development Fund under Grant Number 13/RC/2073 P2. RD acknowledges the funding support from the National Breast Cancer Research Institute grant number FY24001. DG, KF and WT thank the University of Strathclyde for financial support and the EPSRC for providing the instrumentation via EP/N010914/1.

## AUTHOR CONTRIBUTIONS

Conceptualization, PF, NM, RD, AP; methodology, PF, NM, RD, DG, KF, WT, HS, YD, AP; Investigation, NM, WT, YD; writing—original draft, PF, NM; writing—review & editing, PF, NM, WT, AP, RD, DG, KF, HS, YD; funding acquisition, PF, RD, DG, KF, HS, AP; resources, PF, RD, DG, KF, HS, AP; supervision, PF, AP, DG, WT, YD, HS, RD.

## DECLARATION OF INTERESTS

The authors declare no competing interests.

## SUPPLEMENTAL INFORMATION

Document S1. Figures S1, S3-S13, Schemes S14, S15 and Table S2

Table S2. Peak assignments for the Raman spectra acquired from metallacarboranes

## MATERIALS AND METHODS

### Cell culture

All cell lines were acquired from the American Type Culture Collection (ATCC). HCC-1954 were cultured in in Roswell Park Memorial Institute (RPMI 1640) supplemented with 10% foetal bovine serum (FBS, Gibco™, Fisher Scientific), 1% penicillin/streptomycin (Gibco™, 10 000 U/mL, Fisher Scientific) incubation at 37 °C. HDF cell line were grown in Dulbecco’s Modified Eagle Medium (DMEM, glucose: 1 g/L) supplemented with 10% foetal bovine serum (FBS, Gibco™, Fisher Scientific), 1% penicillin/streptomycin (Gibco™, 10 000 U/mL, Fisher Scientific) 5 % CO_2_ and 37 °C. MDA-MB-231 in L-15 GLUTAMAX^R^ supplemented media w/ 10% FBS and 1% PenStrep, 0.5 % CO_2_. For media effect experiments MDA-MB-231 cells were incubated at 5 % CO_2_ and 37 °C for three months in complete DMEM media w/ 10% FBS and 1% PenStrep, and a parallel cell group in complete L-15 GLUTAMAX^R^ supplemented media w/ 10% FBS and 1% PenStrep, 0.5 % CO_2_. All cells were split during the logarithmic phase and fed in three-day intervals.

### Zebrafish maintenance and studies

Fli:GFP casper fish were maintained in tanks in a continuous flow system (Company name) with daily water exchange at 28 °C under a light/dark cycle of 14 h light/10 h dark. The day before mating, the female and male fish were placed in single-cross spawning boxes and separated. The next morning, females and males were placed together to mate, and embryos were collected after 30 min. These were then transferred to Petri dishes (60 larvae per Petri dish) containing egg water and incubated at 28 °C. Petri dishes were checked for dead embryos and water was exchanged daily.

CM-Dil lipophilic dye was diluted according to the manufacturer’s instructions in DMSO at a concentration of 1.0 mg/mL. The concentration was made to 5.0 μg/mL in Dulbecco’s phosphate-buffered saline (DPBS) fresh on the day of cell inoculation. The MDA-MB-231 cells were then trypsinised, counted, and diluted to 1 million cells/mL with complete media. The single-cell suspension was centrifuged in a 15.0 mL falcon tube for five minutes at 200 × g (Eppendorf 5702). The complete medium was removed and the pellet was resuspended in 5.0 mL of DPBS. The single-cell suspension was centrifuged in a 15.0 mL falcon tube for five minutes at 200 × g (Eppendorf 5702), and the supernatant was removed. The pellet was resuspended in 1.0 mL of DPBS and transferred to a 1.5 mL Eppendorf tube. This was then centrifuged for five minutes at 135 × g (Eppendorf 5424), and supernatant removed, followed by resuspension in 5.0–10.0 μL 2 % Polyvinylpyrrolidone (PVP) 40 (helps prevent clumping) with a final cell density of 200– 500 cells/nL. The pellet was resuspended in 1.0 mL CM-Dil solution and transferred to a 1.5 mL Eppendorf tube. The mixture was first incubated for five minutes at 37 °C and then for 15 min at 4 °C. This was then centrifuged for five minutes at 135 × g (Eppendorf 5424), the supernatant was removed, and the pellet was resuspended in 1.0 mL of DPBS and centrifuged again. Removal of the supernatant helped to remove residual CM-Dil that was not internalised by the cells. The cells were resuspended in 2 % PVP40 to prepare a 1 million cells/mL suspension. The cell suspension was maintained at room temperature until the end of injection.

Fluorescence microscopy imaging of the zebrafish was carried out at 0-5 h and 72-75 h post injection of dyed MDA-MB-231 cells with a Leica MZ16FA equipped with a Leica DFC 420 C digital color camera at 475/509 nm excitation/emission for GFP images and 553/570 nm excitation/emission for CM-Dil images. Fluorescence was quantified using ImageJ software.

## METHOD DETAILS

### Cytotoxicity studies

The MTS assay^[50]^ (Abcam) was performed as follows: HCC-1954 and MDA-MB-231 cells were seeded at densities of 5,000 and 10,000 cells per well, respectively, in 96-well plates. These were then left to attach overnight at 37 °C to the respective culture media. This medium was then replaced with 100 μL of relevant drug dilutions in culture media and incubated for the stated treatment periods. This was then replaced by 100 μL of fresh media, to which 10 μL MTS assay which had been warmed to room temperature, was added and incubated at 37 °C for 3 hours. Absorbance was measured in a Viktor X5 plate reader at 490 nm, as per the manufacturer’s instructions.

### Raman

Raman spectra were acquired on a Renishaw InVia Raman microscope equipped with a 532 nm Nd:YAG laser providing a maximum power of 45 mW using a 1800 l/mm grating, a 633 nm HeNe laser providing a maximum power of 17 mW using a 1200 l/mm grating, and a 785 nm diode laser providing a maximum power of 300 mW using a 1200 l/mm grating.

#### Neat samples

A microgram of solid material was transferred to a CaF_2_ disc and imaged directly using the imaging conditions outlined in the corresponding Figure legend.

#### Cell samples

MDA-MB-231 and HCC-1954 cells were plated directly onto a 35 mm Ibidi™ dish with a polymer coverslip at a concentration of 2.5×10^5^ cells/mL in DMEM and RPMI media, respectively. Cells were cultured for 24 h prior to treatment with metallacarboranes (500 μM, 30 min). Raman spectra were acquired using 532 nm excitation with a 60× lens (36 mW) for 5 s.

#### Data processing for Raman spectra

All spectra were processed using WiRE 4.4™ software, enabling cosmic ray removal and baseline subtraction. Peak normalisation was performed in OriginPro2018 software, and the peak areas determined using the Integrate tool.

### SRS microscopy

An integrated laser system (picoEmerald™ S, Applied Physics & Electronics, Inc.) was used to produce two synchronised laser beams at 80 MHz repetition rate. A fundamental Stokes beam (1031.4 nm, 2 ps pulse width) was intensity-modulated by an electro-optic modulator (EoM) with >90% modulation depth, and a tunable pump beam (700–960 nm, 2 ps pulse width, <1 nm (10 cm^−1^) spectral bandwidth) was produced by a built-in optical parametric oscillator. The pump and Stokes beams were spatially and temporally overlapped using two dichroic mirrors and a delay stage inside the laser system, and coupled into an inverted laser-scanning microscope (Leica TCS SP8, Leica Microsystems) with optimised near-IR throughput. SRS images were acquired using a 40× objective lens (HC PL IRAPO 40×, NA). 1.10 water immersion lens) with a 9.75-48 μs pixel dwell time over a 512 × 512 or 1024 × 1024 frame. The Stokes beam was modulated using a 20 MHz EoM. Forward-scattered light was collected using an S1 N. A. 1.4 condenser lens (Leica Microsystems). Images were acquired at a 12-bit image depth. The laser powers measured after the objective lens were in the ranges of 10–30 mW for the pump beam only, 10–50 mW for the Stokes beam only, and 20–70 mW for (pump and Stokes beams). The spatial resolution of the system was approximately 450 nm (pump wavelength = 792 nm).

#### SRS imaging

MDA-MB-231 and HCC-1954 cells were plated on high-precision glass coverslips (#1.5H Thickness, 22 × 22 mm, Thorlabs) in a 6-well plate in DMEM and RPMI 1640 media, respectively, at a concentration of 1.5 × 10^5^ cells/mL and incubated at 37 °C and 5% CO_2_ for 24 h prior to treatment. Cells were treated with metallacarboranes from stocks dissolved in the relevant complete medium and incubated at 37 °C and 5% CO_2_ for the indicated time. The coverslips were then affixed using a nitrocellulose solution in butyl acetone to the glass microscope slides with a PBS boundary between the glass layers prior to imaging. False colour assignments, scale bars, and image overlays were added to the images using ImageJ software. Consistent brightness and contrast settings were used to compare image datasets. Additional information is provided in the SI.

### Data analysis of SRS images

False colour assignments, scale bars and image overlays were added to images using ImageJ software. Consistent brightness and contrast settings were used when comparing image datasets.

### Experimental: synthetic procedures

^1^H, ^11^B and ^13^C NMR spectroscopy was performed on a Varian 500 MHZ 54 mm AR spectrometer. The spectra for ^1^H, ^13^C, and ^11^B were recorded at 500 MHz, 125 MHz and 160.4 MHz respectively. Trimethylsilane was used as a reference for ^1^H and ^13^C, with boron trifluoride diethyl etherate used for ^11^B. Deuterated solvents used included D_2_O, C_2_D_3_N. Solvent stated on spectra in supporting information.

Experiments were carried out, except when noted, under a dry, oxygen-free dinitrogen atmosphere using standard Schlenk techniques, with some subsequent manipulation in the open laboratory. *o*-C_2_B_10_H_12_ was purchased from Katchem and was used as received. All other solvents and organic and inorganic salts were purchased from Across Organics, TCI or Sigma-Aldrich at analytical reagent grade and were used as received. Anhydrous THF was obtained by passing HPLC grade THF through a Grubbs system under inert atmosphere.

Mass spectra were attained using an Agilent 6510 QTOF LC/MS/MS system. Ionization was achieved by electrospray ionization in negative ion mode (ESI−). The capillary voltage was set to 2.5 kV. The cone temperature was 300 °C and the source temperature was 350 °C.

### Preparation of *nido*-carborane

To a solution of 85% KOH (2200 mg, 33.3 mM) in degassed EtOH (14 mL) was added *o*-carborane (1 g, 6.9 mmol) under nitrogen atmosphere. The solution was heated to reflux at 84°C for 2 h. After cooling to room temperature, the solvent was removed under reduced pressure and the solid residue was taken up in distilled water (5 mL). The solution was neutralized with 1 M HCl. Afterwards, an aqueous solution of [HNMe_3_]^+^Cl^-^ was added dropwise until no more precipitate was formed. The white solid was filtered and rinsed with water obtaining [NEt_3_H]^+^ [C_2_B_9_H_12_]^-^ (1.32 g, 6.83 mmol 99%). ^11^B NMR matched signals described in the literature.^[51]^

### Preparation of cobalt (III) bis-dicarbollide Na^+^[CoSAN]

CoSAN was prepared following a procedure reported in the literature, as shown in S15.^[52]^ A solution of NMe_3_H^+^[C_2_B_9_H_12_]^-^ (100 mg, 0.52 µmol) in an aqueous 40% NaOH solution (1 mL) was purged with nitrogen to remove released trimethylamine. To this of CoCl_2_ (109.7 mg, 0.845 µmol) in an aqueous 40% NaOH solution (1 mL) was added, and the solution heated to reflux at 110 °C for 15 mins. This was then diluted with distilled water (5 mL), allowed to cool to room temperature and filtered. Extractions were carried out with filtrate/diethyl ether (3 × 10 mL) and the organic layer retained. The solvent was evaporated, and the residue taken up in distilled water (4 mL). To this a concentrated solution of tetramethylammonium bromide was added, yielding a yellow precipitate. This was filtered, dried *in vacuo* to give NMe_4_^+^[CoSAN]^-^ (72.41 mg, 182 µmol, 70%). NMR spectra matched signals described in the literature as seen in Figure S10, S11.^[52]^ The sodium salt prepared as described in the literature.^[53]^

### Preparation of iron (III) bis-dicarbollide Na^+^[FeSAN]

FeSAN was prepared following a procedure reported in the literature, as shown in Figure S15.^[52]^ A solution of NMe_3_H^+^[C_2_B_9_H_12_]^-^ (100 mg, 0.52 µmol) in an aqueous 40% NaOH solution (1 mL) was purged with nitrogen to remove released trimethylamine. To this FeCl_2_•4H_2_O (218.69 mg,1.11 mmol) in an aqueous 40% NaOH solution (1 mL) was added, and the solution heated to reflux at 110 °C for 15 mins. This was then diluted with distilled water (5 mL), allowed to cool to room temperature and filtered. Extractions were carried out with filtrate/diethyl ether (3 × 10 mL) and the organic layer retained. The solvent was evaporated, and the residue taken up in distilled water (4 mL). To this a concentrated solution of tetramethylammonium bromide was added, yielding a black-red precipitate. This was filtered, dried *in vacuo* to give NMe_4_^+^[FeSAN]^-^ (48.25 mg, 122.2 µmol, 47%), ^11^B NMR matched signals described in the literature as seen in Figure S12.^[52]^ The sodium salt was prepared as described in the literature.^[53]^

### Preparation of nickel (III) bis-dicarbollide Na^+^[NiSAN]

NiSAN was prepared following a procedure reported in the literature, as shown in Figure S15.^[52]^ A solution of NMe_3_H^+^[C_2_B_9_H_12_]^-^ (100 mg, 0.52 µmol) prepared as described in the literature in an aqueous 40% NaOH solution (1 mL) was purged with nitrogen to remove released trimethylamine. To this a NiCl_2•_6H_2_O (109.17 mg, 0.84 µmol) solution in 40% NaOH (1 mL) was added, and the solution heated to reflux at 110 °C for 15 mins. This was then diluted with distilled water (5 mL), allowed to cool to room temperature and filtered. Extractions were carried out with filtrate/diethyl ether (3 × 10 mL) and the organic layer retained. The solvent was evaporated, and the residue taken up in distilled water (4 mL). To this a concentrated solution of tetramethylammonium bromide was added, yielding a yellow-orange precipitate. This was filtered and dried *in vacuo* to give NMe_4_^+^[NiSAN]^-^ (66.13 mg, 166.4 µmol, 64%), ^11^B NMR matched signals described in the literature as seen in Figure S13, as did the mass spectrum shown in Figure S14.^[52]^ The sodium salt was prepared as described in the literature.^[53]^

### Preparation of chromium (III) bis-dicarbollide Na^+^[CrSAN]

CrSAN was prepared following a procedure reported in the literature as seen in Figure S17.^[54]^ CrCl_3_ (39.73 mg, 0.25 mmol) in anhydrous THF (0.5 mL) was added via degassed syringe to a solution of 2[Na]^+^[C_2_B_9_H_11_]^2-^ (88.6mg, 0.5 mmol) (prepared as described in the literature)^[52]^ in anhydrous THF (0.5 mL). This was heated at 72 °C to reflux for 3 h under inert atmosphere. The solvent was evaporated under reduced pressure. Extractions were carried out with filtrate/diethyl ether (3 × 10 mL) and the organic layer retained. The solvent was evaporated, and the residue taken up in distilled water (4 mL). To this a concentrated solution of tetramethylammonium bromide was added, yielding a purple precipitate. This was filtered, washed with distilled water and dried *in vacuo* to give NMe_4_^+^[CrSAN]^-^ (62 mg, 158.6 µmol, 61%), ^11^B NMR matched signals described in the literature as shown in Figure S16.^[54]^ The sodium salt was prepared as described in the literature.^[53]^

### Isolation of sodium metallacarborane salts

Following a reported procedure Amberlite® IR-120 strongly acidic cation exchange resin was used to generate sodium salts of metallacarboranes.^[53]^ Approximately 2/3 of a column was filled with the resin beads, and they were left in 3 M HCl overnight. It was then washed with 250 mL 3 M HCl and washed with deionised water until a neutral pH was obtained. Then, a 3 M NaCl solution is passed through the column until the eluent no longer has an acidic pH. The column is then washed with water until the Tollen’s test indicates the column is free of salts. A solution of the desired [NMe_4_] metallacarborane salt is dissolved in 50:50 ACN:H_2_O and then passed through the column three times. This is evaporated to dryness and extracted with brine/diethyl ether to remove salts. The organic solvent was removed under reduced pressure, the precipitate washed with cyclohexane and dried under vacuum.

## QUANTIFICATION AND STATISTICAL ANALYSIS

### Cell viability

All statistical analysis of cell viability data was done using the t-test function on excel to generate p-values, with averages of technical and experimental replicates.

### ROS generation

All statistical analysis of reactive oxygen species data was done using the t-test function on excel to generate p-values, with averages of technical and experimental replicates.

### Uptake via SRS

All statistical analysis of cell viability data was done using the t-test function on excel to generate p-values, with averages of technical and experimental replicates.

## Notes

### Competing Interest Statement

The authors have declared no competing interest.

