## Supplementary Information for "Unprecedented Cytotoxicity of Transition Metal Metallacarboranes on Triple Negative Breast Cancer Cells and a Vertebrate Cancer Model"

### Contents

**Figures S1-S16**……..…………………..……………………………………….…………………3

**Table S1**………………………………………………………………………….………….……  11

**Schemes S15, S17**……………………………………………………………………….………12

**References**……………………………………………….……………..…………………..…12

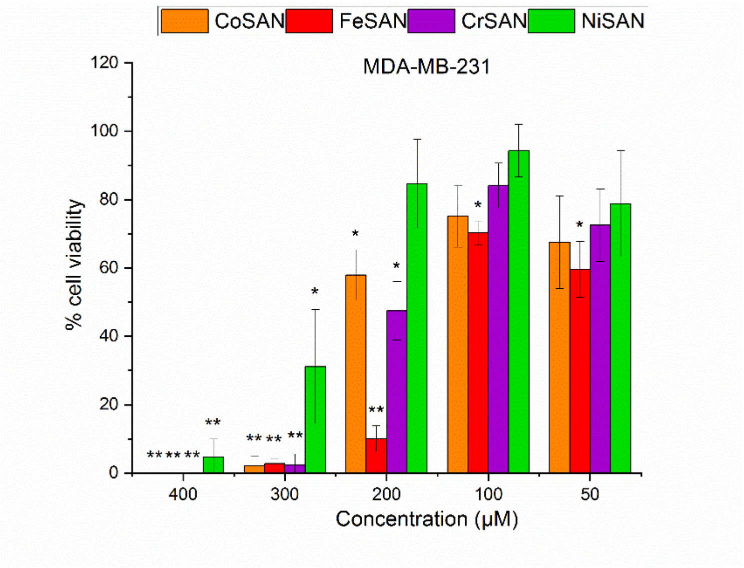

**Figure S1** Cell viability of MDA-MB-231 cell line after 24-hour treatment with stated concentrations of metallacarboranes. *P ≤ 0.05 **P ≤ 0.01 ***P ≤ 0.001 (student's t-test).

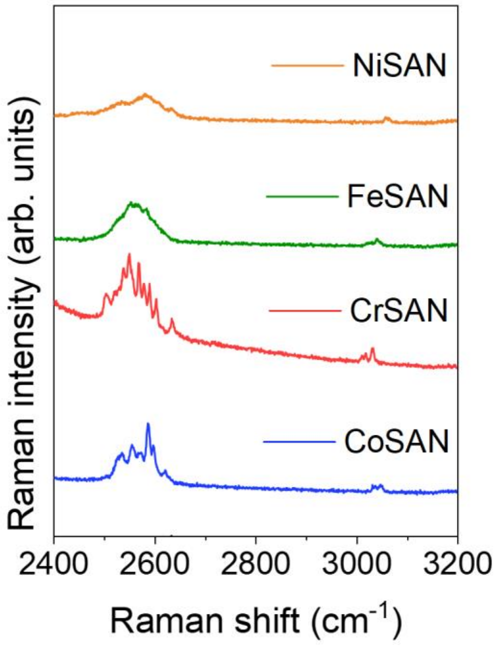

**Figure S2**. Raman spectra of the metallacarboranes acquired in solid form using 785 nm excitation with a 20× lens (~20 mW) for 10 s. The spectra are normalised (0-1) and offset for clarity. Peak assignments are in $cm^{-1}$. [1]

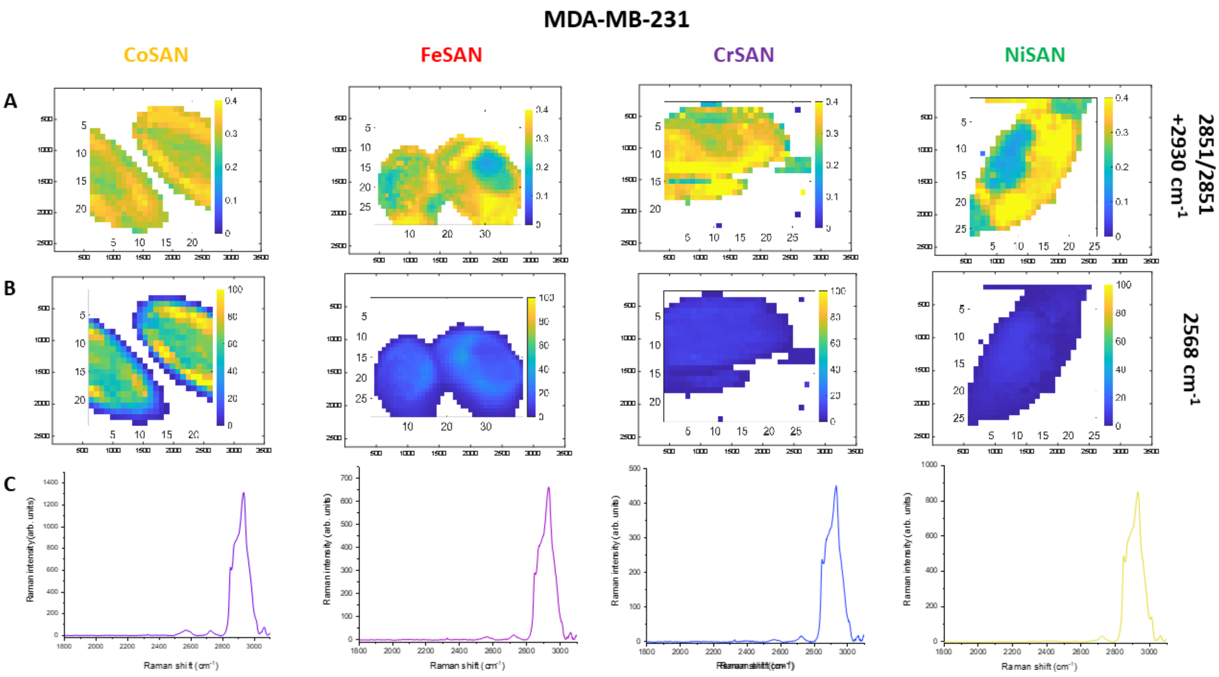

**Figure S3** Spontaneous Raman mapping of MDA-MB-231 cells. Cells were treated with 500 µM of the metallacarboranes in DMEM then mapping of a cell carried out at 2851, 2930 and 2568 $cm^{-1}$ to determine presence of B–H stretching **A**,**B**. Single acquired spectra of the cellular environment is shown in **C**, showing high signal at around 2930 $cm^{-1}$ and comparably weaker signals in the 2568 $cm^{-1}$ region.

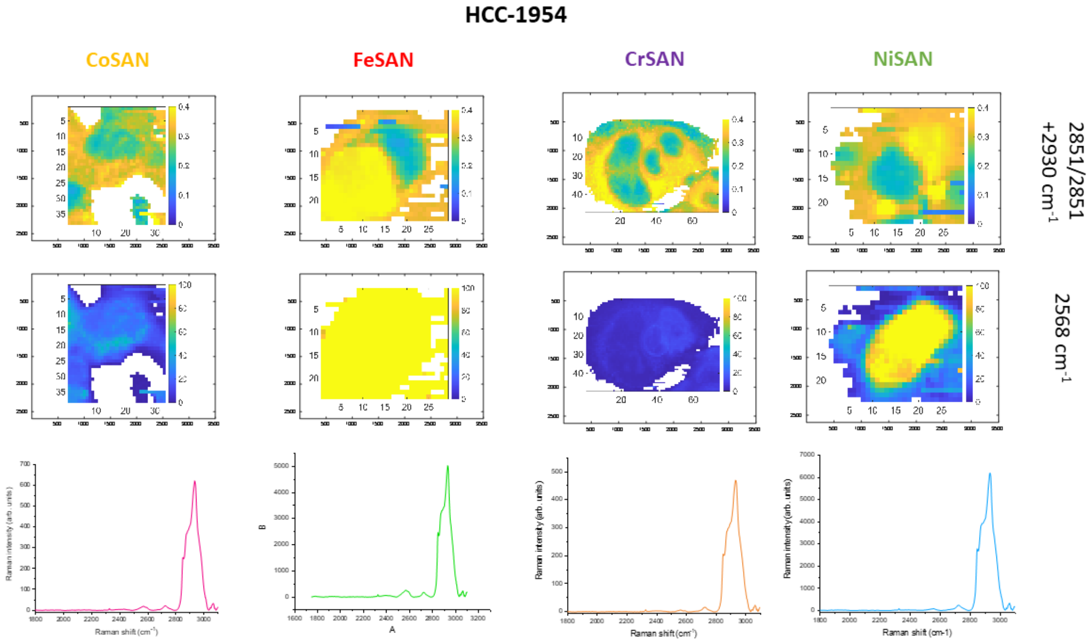

**Figure S4** Spontaneous Raman mapping of HCC-1954 cells. Cells were treated with 500 µM of the metallacarboranes in RPMI for 15 min then mapping of a cell carried out at 2851, 2930 and 2568 $cm^{-1}$ to determine presence of B–H stretching **A**,**B**. Single acquired spectra of the cellular environment is shown in **C**, showing high signal at around 2930 $cm^{-1}$ and comparably weaker signals in the 2568 $cm^{-1}$ region.

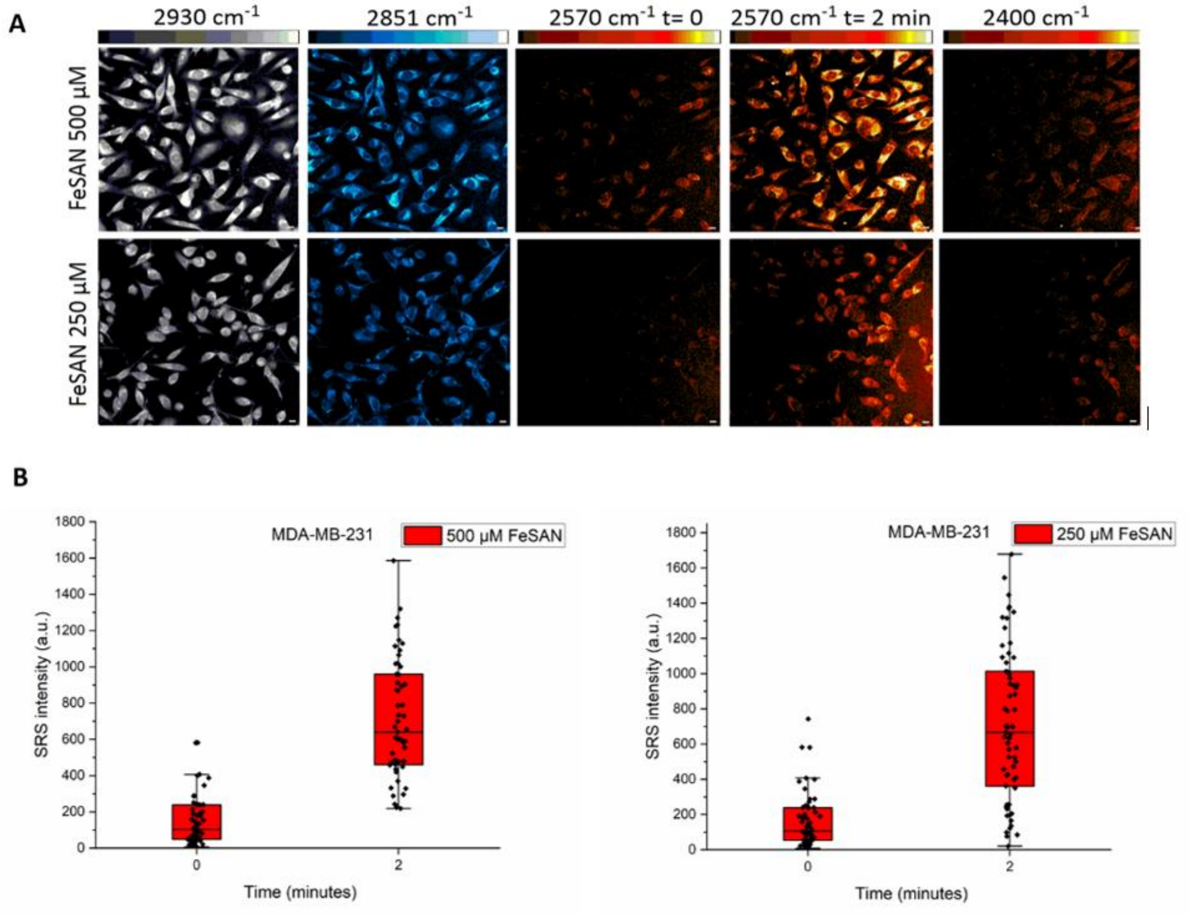

**Figure S5** Uptake imaging of FeSAN. **A** SRS images of MDA-MB-231 cells were acquired at the following frequencies: 2930 $cm^{-1}$ ($CH_3$, proteins), 2851 $cm^{-1}$ ($CH_2$, lipids), 2570 $cm^{-1}$ (B–H) and 2400 $cm^{-1}$ (off-resonance), then treated with FeSAN (500, 250 µM) and imaged at 2570 $cm^{-1}$ (B–H) after approximately 2 minutes. Scale bars: 10 µm. **B** SRS intensities of cells at 2570 $cm^{-1}$ (B–H) before and after treatment.

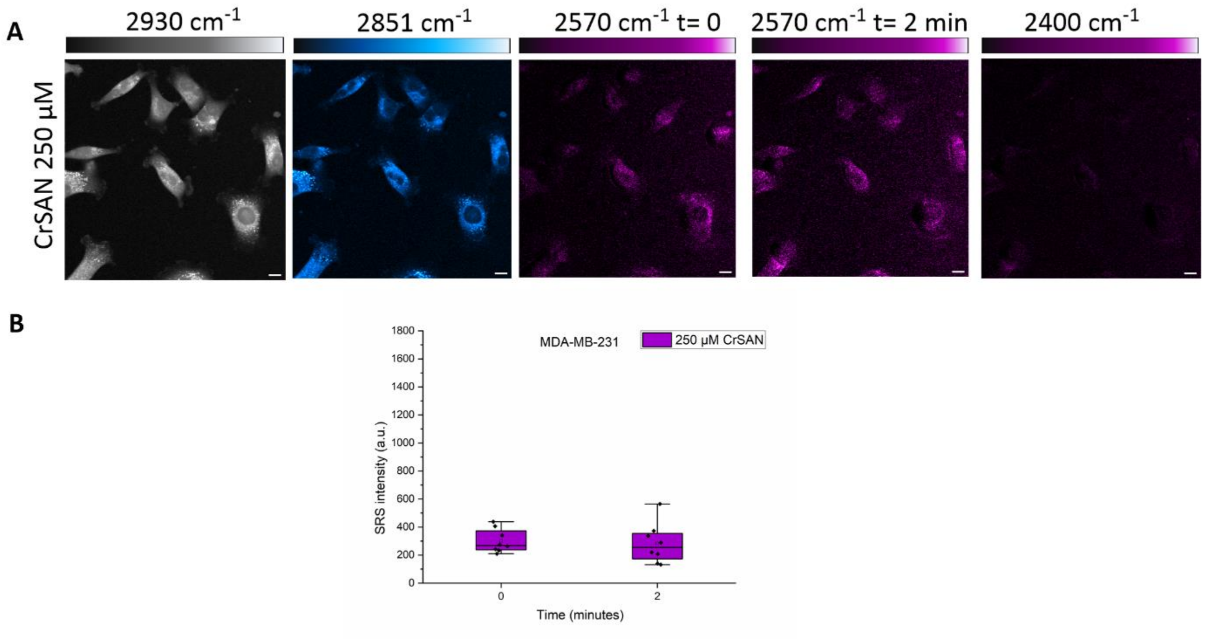

**Figure S6** Uptake imaging of CrSAN **A** SRS images of MDA-MB-231 cells were acquired at the following frequencies: 2930 cm$^{-1}$ ($CH_3$, proteins), 2851 cm$^{-1}$ ($CH_2$, lipids), 2570 cm$^{-1}$ (B–H) and 2400 cm$^{-1}$ (off-resonance), then treated with CrSAN (250 µM) and imaged at 2570 cm$^{-1}$ (B–H) after approximately 2 minutes. Scale bars: 10 µm. **B** SRS intensities of cells at 2570 cm$^{-1}$ (B–H) before and after treatment.

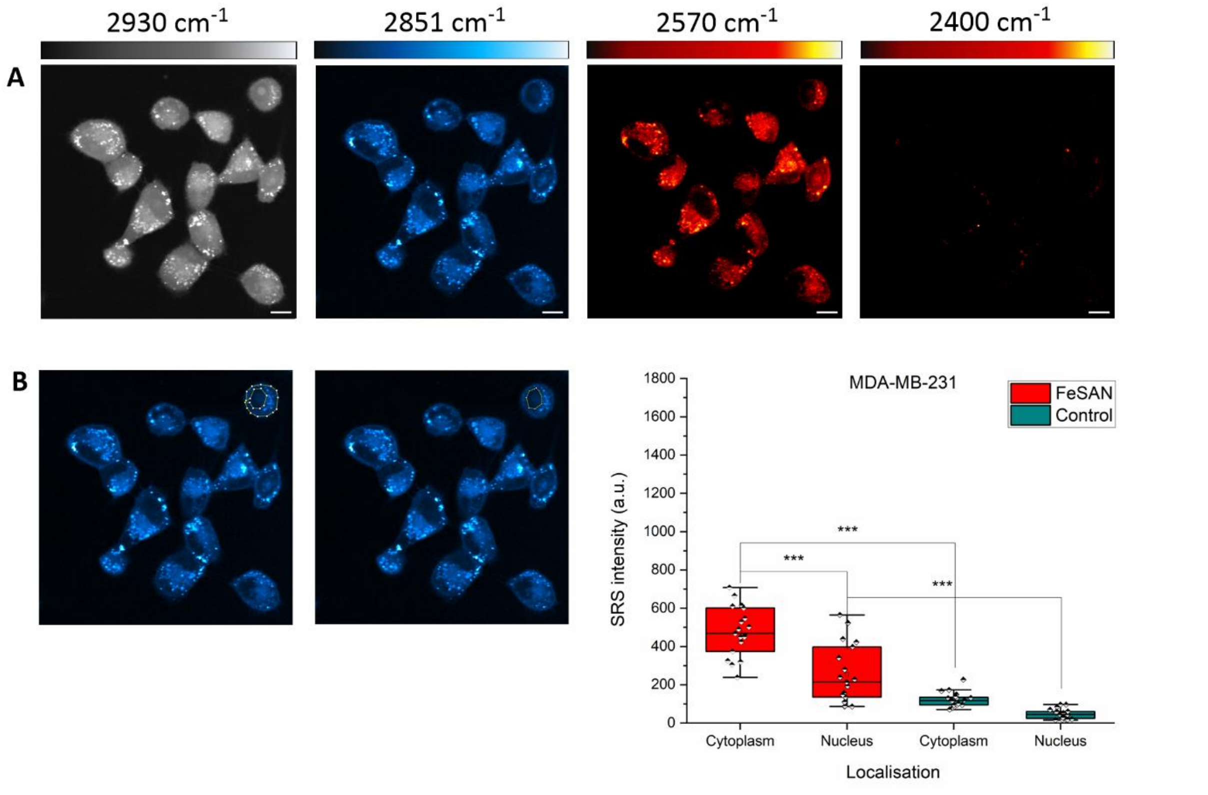

**Figure S7** Cellular localisation of FeSAN within MDA-MB-231 cell line. **A** MDA-MB-231 cells were treated with FeSAN (500 µM, 15 minutes) and SRS images of cells were acquired at the following frequencies: 2930 cm$^{-1}$ ($CH_3$, proteins), 2851 cm$^{-1}$ ($CH_2$, lipids), 2570 cm$^{-1}$ (B–H) and 2400 cm$^{-1}$ (off-resonance. Scale bars: 10 µm. **B** Determination of SRS intensity of 2570 cm$^{-1}$ (B–H) signal in nucleus and cytoplasm.

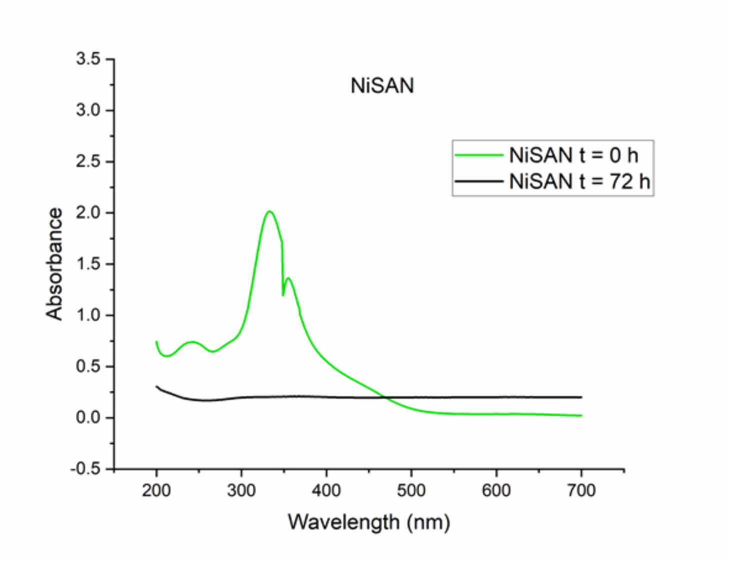

**Figure S8** UV-Vis spectrum of NiSAN over 72 h in PBS at 37 °C.

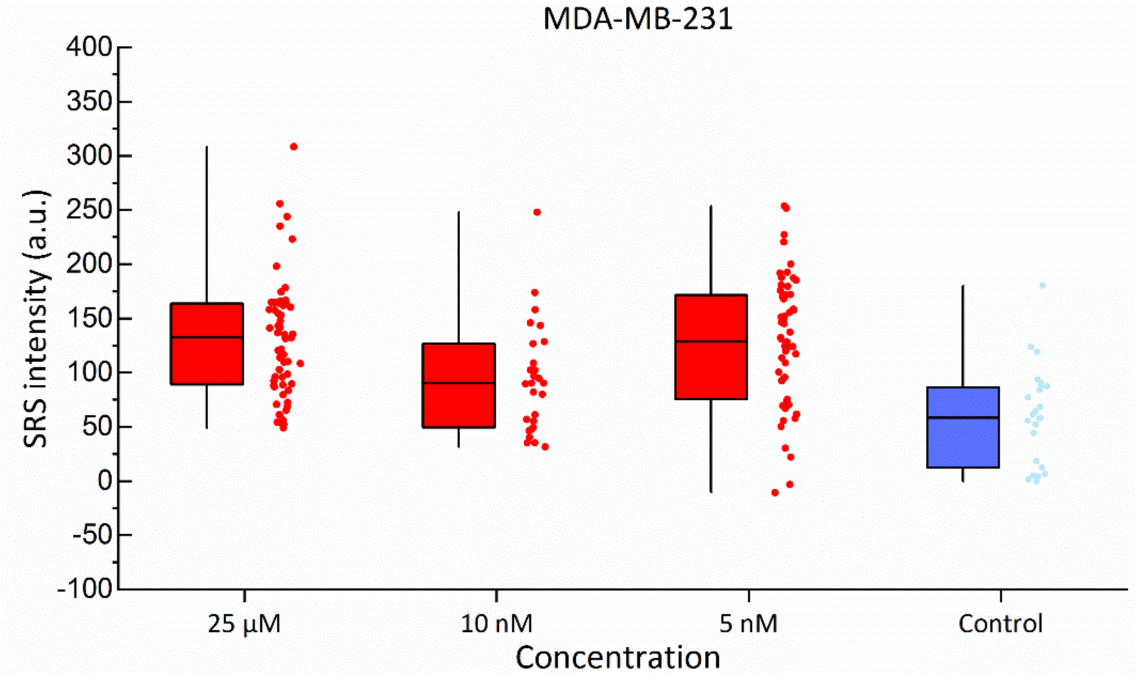

**Figure S9** SRS intensities of B–H stretch at 2570 $cm^{-1}$ within MDA-MB-231 cells following incubation with stated concentrations of FeSAN compared to control.

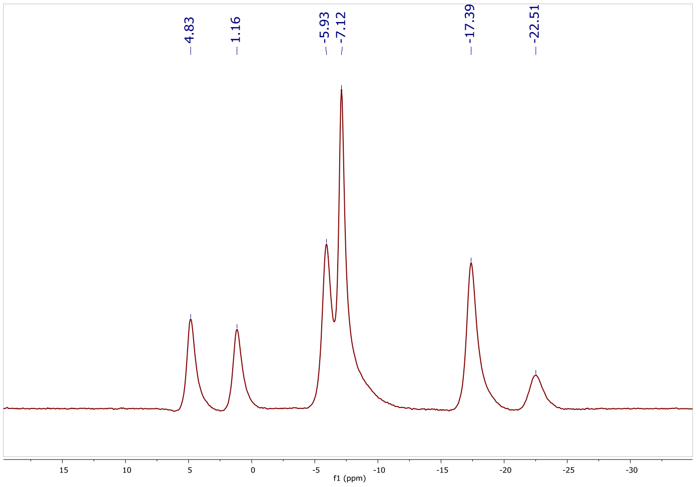

**Figure S10** $^{11}B\{^{1}H\}$ NMR spectrum of $Na^{+}$[CoSAN] in $D_2O$.

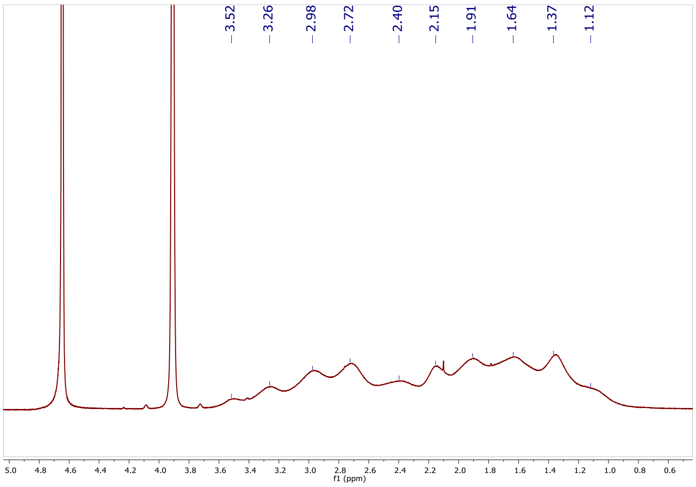

**Figure S11** $^{1}H$ NMR spectrum of $Na^{+}$[CoSAN] in $D_2O$.

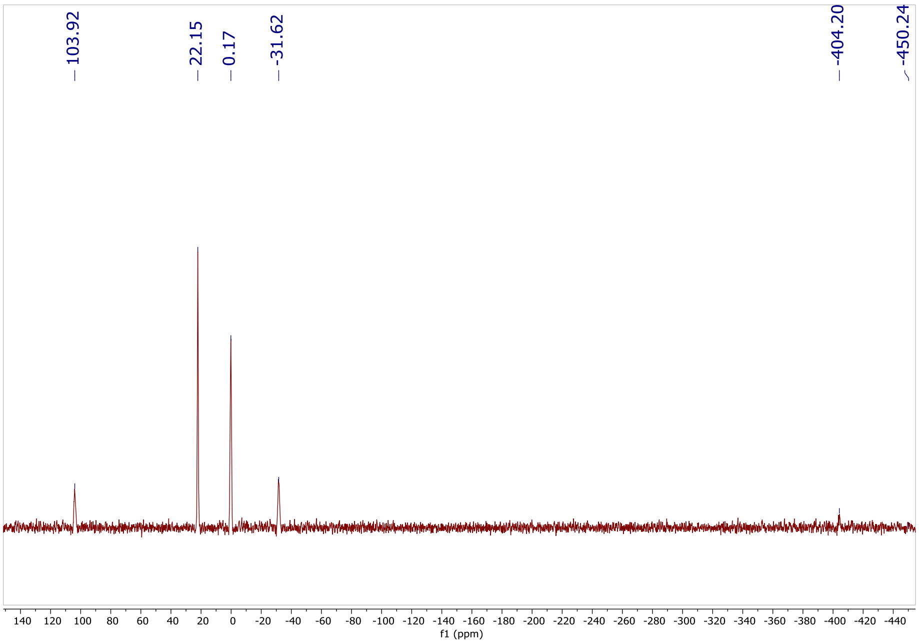

**Figure S12** $^{11}B\{^{1}H\}$ NMR spectrum of $Na^{+}$[FeSAN] in $D_2O$.

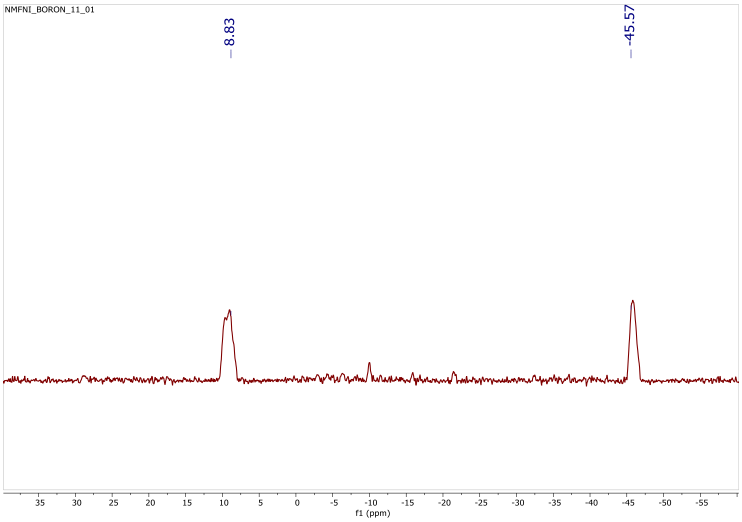

**Figure S13** $^{11}B\{^{1}H\}$ NMR spectrum of $Na^{+}$[NiSAN] in $D_2O$.

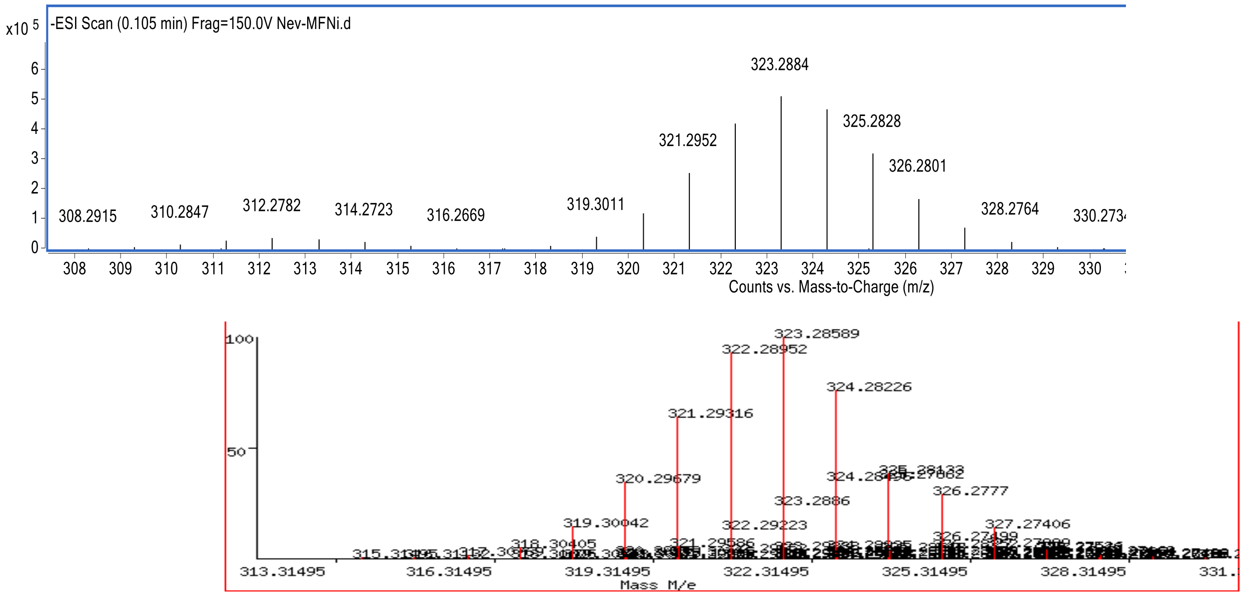

**Figure S14** Mass spectrum of [Na][NISAN] (top) matching predicted distribution (bottom) (323.2884 m/z).

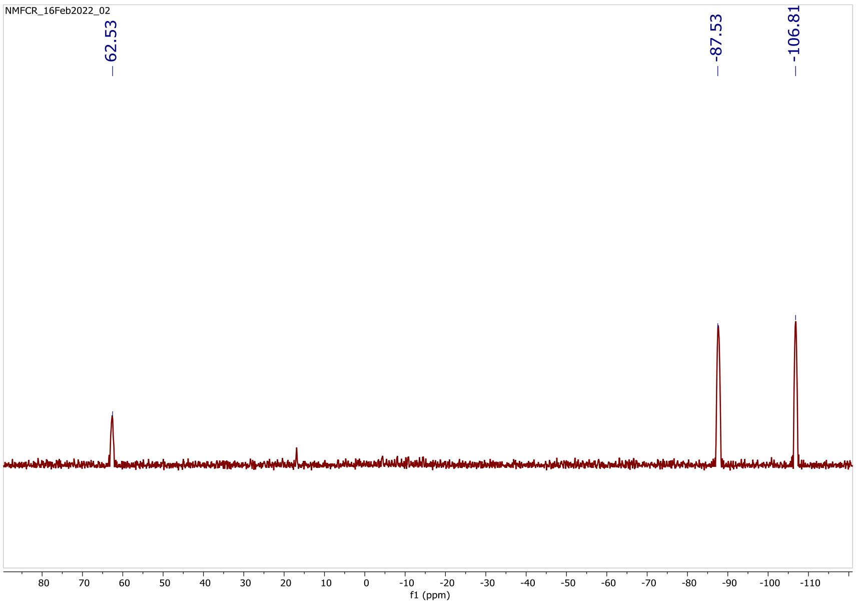

**Figure S16** $^{11}B\{^{1}H\}$ NMR spectrum of $Na^{+}$[CrSAN] in $D_2O$.

**Table S1** Peak assignments for the Raman spectra acquired from metallacarboranes (M = Co, Cr, Fe, Ni).[1]

| Raman shift ($cm^{-1}$) | | | | |
|---|---|---|---|---|
| **CoSAN** | **CrSAN** | **FeSAN** | **NiSAN** | **Assignment** |
| 209.2 (vs) | 189.7 (vs) | 201.4 (vs) | 168.9 (vs) | Whole molecule stretching[a] |
| 276.1 (s) | 238.9 (s) | 242.8 (s) | 224.7 (s) | Carborane cages rocking[a] |
| 586.0 | 574.1 | 562.2, 580.2 | 559.5, 614.0 | |
| 635.7 | 632.1 | 633.3 | 638.1 | B-B-M bending[a] |
| | | | 687.1 | |
| 728.6 | 729.8 | 728.6 | | |
| 753.3 | 742.7 | 751.0 | 739.2 | Icosahedral breathing mode[a] |
| 793.1 | 779.1 | 786.1 | , 793.1 | |
| 875.3 | 881.1 | | 840.8, 862.7 | |
| 917.7 | 915.5 | 918.9 | 916.6 | |
| 985.7 | 988.0 | 981.2 | 983.5 | |
| 1005.7 | 1008.2 | 1001.5, | 995.9 | |
| 1019.4 | 1022.8 | 1020.5 | 1023.9 | |
| 1142.2 | 1152.0 | 1145.5 | 1147.7, 1227.9 | δ(CH)[b] |
| 2535.3, 2554.9, 2572.8, 2586.6, 2598.7, 2621.2 | 2503.3, 2537.8, 2550.0, 2567.9, 2589.8, 2603.5, 2634.0 | 2552.7, 2568.9, 2582.6, | 2534.5, 2580.1, 2609.1, 2634.8 | B–H stretching[a,b] |
| 3031.9, 3042.1, 3047.9 | 3010.8, 3018.8, 3033.4 | 3022.1, 3030.6, 3042.6 | 3057.3, 3063.8 | C-H stretching[a,b] |

Notes: where no assignment has been made, Raman spectral frequencies have been grouped based on similar wavenumber positions. Most vibrations having frequencies in the region from 450 to 1100 $cm^{-1}$ are of complex origin, and therefore challenges remain over their interpretation and assignment. References: [a] B. Barszcz *et al. J. Mol. Struct.*, **2010**, *976*, 196-199; [b] L. A. Leites, *Chem. Rev.*, **1992**, *92*, 279-323. (vs) very strong, (s) strong.

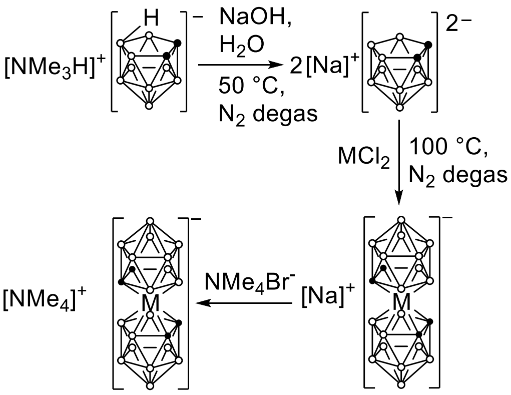

**Scheme S15** Preparation of Co, Fe, Ni metallacarboranes.[2]

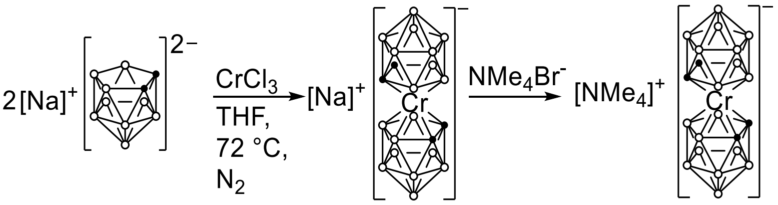

**Scheme S17** Procedure for synthesis of Cr metallacarborane.[3]
